# AI-Driven Spatial Transcriptomics Unlocks Large-Scale Breast Cancer Biomarker Discovery from Histopathology

**DOI:** 10.1101/2024.10.16.618609

**Authors:** Eldad D. Shulman, Emma M. Campagnolo, Roshan Lodha, Amos Stemmer, Thomas Cantore, Beibei Ru, Tian-Gen Chang, Sumona Biswas, Saugato Rahman Dhruba, Andrew Wang, Rohit Paul, Sarath Kalisetty, Tom Hu, Maclean Nasrallah, Sheila Rajagopal, Stephen-John Sammut, Stanley Lipkowitz, Peng Jiang, Carlos Caldas, Simon Knott, Danh-Tai Hoang, Kenneth Aldape, Eytan Ruppin

**Affiliations:** Cancer Data Science Laboratory, Center for Cancer Research, National Cancer Institute, Bethesda, MD, USA; Faculty of Medical and Health Sciences, Tel Aviv University, Tel Aviv, Israel; Office of the Director, Center for Cancer Research, National Cancer Institute, National Institutes of Health, Bethesda, MD, USA; Division of Neuropathology, Department of Pathology and Laboratory Medicine, Perelman School of Medicine, University of Pennsylvania, Philadelphia, PA, USA; Women’s Malignancies Branch, Center for Cancer Research, National Cancer Institute, Bethesda, MD, USA; Breast Cancer Now Toby Robins Research Centre, Institute of Cancer Research, London, UK; The Royal Marsden Hospital NHS Foundation Trust, London, UK; Department of Clinical Biochemistry and Institute of Metabolic Science, University of Cambridge, UK; Department of Biomedical Sciences, Cedars-Sinai Medical Center, Los Angeles, CA, USA; Laboratory of Pathology, Center for Cancer Research, National Cancer Institute, Bethesda, MD, USA

## Abstract

Spatial transcriptomics (ST) is transforming our understanding of tumor heterogeneity by enabling high-resolution, location-specific mapping of gene expression across tumors and their microenvironment. However, the associated high cost of the assay has limited cohort size and hence large-scale biomarker discovery. Here we present *Path2Space*, a deep learning approach that predicts spatial gene expression directly from histopathology slides. Trained on substantial breast cancer ST data, it robustly predicts the spatial expression of over 4,300 genes in independent validations, markedly outperforming existing ST predictors. *Path2Space* additionally accurately infers cell-type abundances in the tumor microenvironment (TME) based on the inferred ST data. Applied to more than a thousand breast tumor histopathology slides from the TCGA, *Path2Space* characterizes their TME on an unprecedented scale and identifies three new spatially-grounded breast cancer subgroups with distinct survival rates. *Path2Space*-inferred TME landscapes enable more accurate predictions of patients’ response to chemotherapy and trastuzumab directly from H&E slides than those obtained by existing established sequencing-based biomarkers. *Path2Space* thus offers a transformative, fast and cost-effective approach to robustly delineate the TME directly from their histopathology slides, facilitating the development of spatially-grounded biomarkers to advance precision oncology.

## Introduction

The rise of spatial transcriptomics (ST) is transforming our understanding of tumor heterogeneity by enabling high-resolution, location-specific mapping of gene expression across tumors and their microenvironments. For example, in lung adenocarcinoma, ST revealed macrophage phenotypic diversity^1,2^. In breast cancer, it illuminated immune-enriched clusters with varying infiltration patterns^3,4^ and revealed recurrent neoplastic cell heterogeneity, leading to the identification of nine ’ecotypes’ with unique cellular compositions and clinical outcomes^5^, and yielded non-spatial tertiary lymphoid structures bulk gene signatures associated with disease outcome and therapy response^6^. In glioblastoma, it uncovered distinct transcriptional programs of sub-clonal structures^7^. It has also characterized the architecture of invasive tumors in melanoma^8^, enabled 3D reconstruction of colorectal cancer^9^, and allowed exploration of intratumoral microbial communities^10^. In HPV-negative oral squamous cell carcinoma, it revealed distinct transcriptional profiles in the tumor core and leading edge, with implications for prognosis and drug response across multiple cancer types^11^. Moreover, spatial transcriptomics has elucidated the intricate cross-talk between cancer cells and proximal immune cells in the tumor microenvironment (TME), providing insights into tumor progression and potential therapeutic targets^12^. By providing unprecedented spatial context to gene expression, spatial transcriptomics holds immense potential to unlock novel biomarkers, decipher treatment resistance mechanisms, and guide more precise cancer therapies.

Despite these remarkable advances, the translational potential of spatial transcriptomics remains constrained by its high cost^13^. The latter hinders the assembly of large patient cohorts, essential for discovering robust biomarkers, developing predictive response models, and ensuring the statistical power needed to validate findings across diverse populations. Given that spatial transcriptomics data is often paired with aligned H&E images, predicting spatial gene expression from H&E slides presents an opportunity to extend these insights at scale. Deep learning applied to H&E images has already shown promise in identifying cell types^14^ and molecular features not apparent to human observers, including genetic mutations^15^, bulk mRNA expression^16^, and methylation patterns^17^.

Previous efforts to predict spatial gene expression from H&E images have shown promise but had limited predictive ability, robustly predicting the expression of only a few hundred genes^18–26^. Given their limited prediction capacity, these studies have not been applied to predict patient survival and treatment response in larger clinical cohorts where only H&E images are available. Expanding our ability to predict spatial transcriptomics features is hence essential for uncovering their biological and clinical significance while avoiding the prohibitive costs of their direct measurement.

To address this challenge, we developed *Path2Space*, a model which predicts spatial gene expression from histopathology slides with unprecedented accuracy focusing on breast cancer due to its prevalence and rich data availability. The model demonstrates robust predictive performance on external datasets and can reliably infer localized cell type compositions from the associated predicted spatial gene expression patterns. When applied to the breast cancer cohort from The Cancer Genome Atlas (TCGA), *Path2Space* identifies three new prognostic breast cancer subgroups based on their predicted spatial transcriptomic cluster compositions, a classification framework we term *SpatioTypes*. Applied to two large cohorts of breast cancer patients treated with trastuzumab and with chemotherapy, *Path2Space* uncovers spatial biomarkers and predicts therapy response at accuracy levels higher or equivalent to those obtained from measured bulk tumor transcriptomics. *Path2Space* is a new, generic approach that forms the basis for similar analyses of large clinical datasets in different indications and treatments, enabling discoveries currently unattainable with the limited scale of experimental existing spatial transcriptomics studies.

## Results

### Study overview

We developed *Path2Space*, a deep-learning model to predict spatial gene expression from H&E slide images of breast cancer tumors, aiming to uncover microenvironment characteristics and other spatial biomarkers associated with treatment response.

To develop the spatial gene expression predictor, we trained a deep learning model on 10X genomics Visium ST data from slides with matched H&E images, where gene expression profiles were derived from 55 μm diameter areas termed spots. The prediction model consists of two main components: preprocessing and regression (**Fig. 1a**; **Methods**). During preprocessing, tile images surrounding individual spots are extracted, color-normalized, and processed using CTransPath, a foundational digital pathology model for feature extraction^27^. In the regression step, a multi-layer perceptron (MLP) neural network is trained to predict the gene expression vector (14,068 genes) for each spot from its corresponding tile image. Following model training, we applied a spatial smoothing technique to mitigate technical variability in spatial transcriptomics data. This approach averages the predicted gene expression values of each spot with its immediate neighbors. The same smoothing was applied to the original expression measurements (**Supplementary Fig. 1a**) and incorporated into model evaluation and downstream analyses. The model was trained using cross-validation on the Bassiouni et al. cohort^4^, comprising 41,887 matched image and gene expression profile spots, which passed image processing quality control (QC, **Methods**), derived from 25 ST tissue sections from 14 breast cancer patients. It was validated on two independent external ST datasets: one cohort consisting of 14,510 QC-passed spots, from four tissue sections from three patients, obtained from 10x Genomics^28^, and another cohort from Janesick et al.^29^, consisting of 4,301 QC-passed spots from a single section from one patient.

**Fig. 1:**
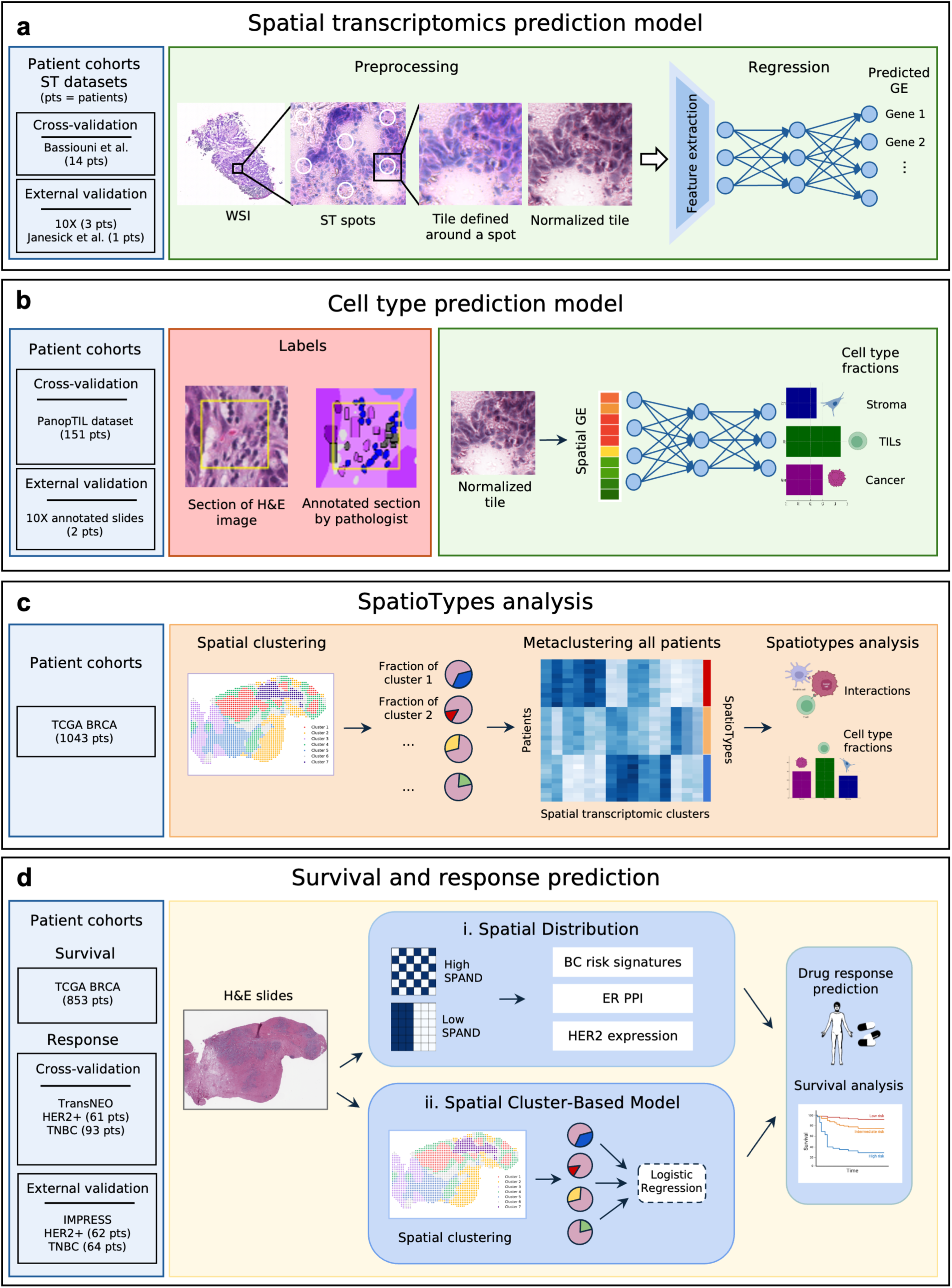
Overview of the datasets and computational workflow. **a**, The *Path2Space* spatial gene expression model was trained and evaluated on three datasets with paired H&E and spatial transcriptomics from the Bassiouni et al. cohort. Whole slide images (WSIs) were partitioned into tiles around each of the spatial transcriptomics spots. These tiles underwent color normalization followed by feature extraction using the CTransPath digital pathology foundational model. The extracted features were then input into a multi-layer perceptron (MLP) to infer gene expression for each tile. **b**, The *Path2Space* cell type fraction estimation model was trained on PanopTILs cell type annotations, using the 10x and Janesick et al. cohorts for external validation (left). The model estimates cell type fractions per tile using an MLP that takes as input the inferred spatial gene expression from whole slide images (right). **c**, *Path2Space* predicted spatial gene expression from TCGA BRCA WSIs, which were used to identify spatial clusters that define tumors spatial profiles. Based on these profiles, an unsupervised algorithm grouped tumors into distinct, prognostic, spatially-grounded breast cancer subgroups, termed *SpatioTypes*. **d,** Two approaches for predicting patient outcomes: (i) <u>Spatial distribution:</u> We defined *SPAtial Neighborhood Diverseness (SPAND)* to measure gene expression spatial patterns, focusing on genes from OncotypeDX and MammaPrint signatures, ER protein-protein interactions (PPI), and HER2. (ii) <u>Spatial Cluster-Based Model</u>: Using spatial cluster compositions from approach (**c**), we trained logistic regression models to predict treatment response, evaluated through cross-validation on TransNEO and further validated on the independent IMPRESS cohort.

The second step of *Path2Space* is to estimate cell type fractions at Visium spot resolution. To this end, we leveraged the PanopTILs dataset^30^, which includes annotations of individual nuclei cell type identities from 1,709 regions of interest (ROIs) across 151 TCGA breast cancer patients. A cell type predictor was trained for the three most prevalent cell types—cancer cells, lymphocytes, and stromal cells. Tile images matching the size of Visium spots (pseudospots) were extracted, and the fractions of each cell type within each image were computed. To train and evaluate the predictor in cross-validation, *Path2Space* was used to generate gene expression for the pseudospot images, which were then used as input to an MLP regressor to predict cell type fractions. External validation was conducted using H&E images with 15,328 spot-level cell type annotations from the 10x cohort (**Fig. 1b**).

Thirdly, our main goal is to apply the trained model (of both ST and cell abundances) to discover *spatial biomarkers* of patient survival and treatment response from the pathology slides of large patient cohorts. To this end, we applied *Path2Space* to the TCGA breast cancer cohort, identifying and characterizing spatial architectures associated with disease-free survival (**Fig. 1c**). We also applied *Path2Space* to study TMEs and infer treatment response biomarkers in two additional HER2-positive (HER2+) breast cancer cohorts with treatment response data: TransNEO^31^ (chemotherapy n=93, trastuzumab n=61) and IMPRESS^32^ (chemotherapy n=64, trastuzumab n=62) (**Fig. 1d**), and benchmarked *Path2Space* prediction performance compared to a broad array of extant biomarkers.

### Predicting spatial gene expression from H&E slides

We trained a *Path2Space* breast cancer model based on the Bassiouni et al. ST dataset, consisting of H&E slides and matched gene expression profiles obtained via the Visium platform. We selected 25 high-quality slides from 14 triple-negative breast cancer (TNBC) patients out of 36 based on spot density (**Methods**). These slides contained an average of 1276 ± 444 spots (mean ± SD) and had 4594 ± 2465 detected genes per spot (expression values>0) (**Supplementary Fig. 2a-c**). Due to the high proportion of missing values, typical of ST data, we focused on 14,068 genes that were detected in ≥5% of spots on each slide.

To evaluate our model, we employed a leave-one-patient-out cross-validation approach. The model was iteratively trained on slides from all except one patient, and then tested on the slides of the held-out patient. We assessed gene-wise prediction accuracy by calculating the average Pearson Correlation Coefficient (PCC) between spatially smoothed (overview) actual and predicted expression values across all spots in each slide (**Methods**). This yielded a distribution of gene-wise correlations with a median of 0.37 (**Fig. 2a**). Notably, 6,022 out of 14,068 genes (43%) demonstrated a mean correlation above 0.4 (**Fig. 2a; Supplementary Fig. 1b**), indicating robust predictive performance for a substantial portion of the transcriptome. In addition to evaluating the ability of *Path2Space* to predict continuous gene expression values, we examined its ability to classify high and low expression states. Binarizing the measured gene expression values within each slide as high or low relative to their median, we observed a median AUC of 0.69 across all genes, with 2,137 genes exceeding an AUC of 0.75 (**Fig. 2b**). We observed a strong correlation between the abundance of specific gene measurements (fewer spots with missing values) and *Path2Space’s* ability to predict their gene expression (**Supplementary Fig. 2d**), suggesting that prediction accuracy could improve in the future with the advent of ST assays of higher sensitivity.

**Fig. 2:**
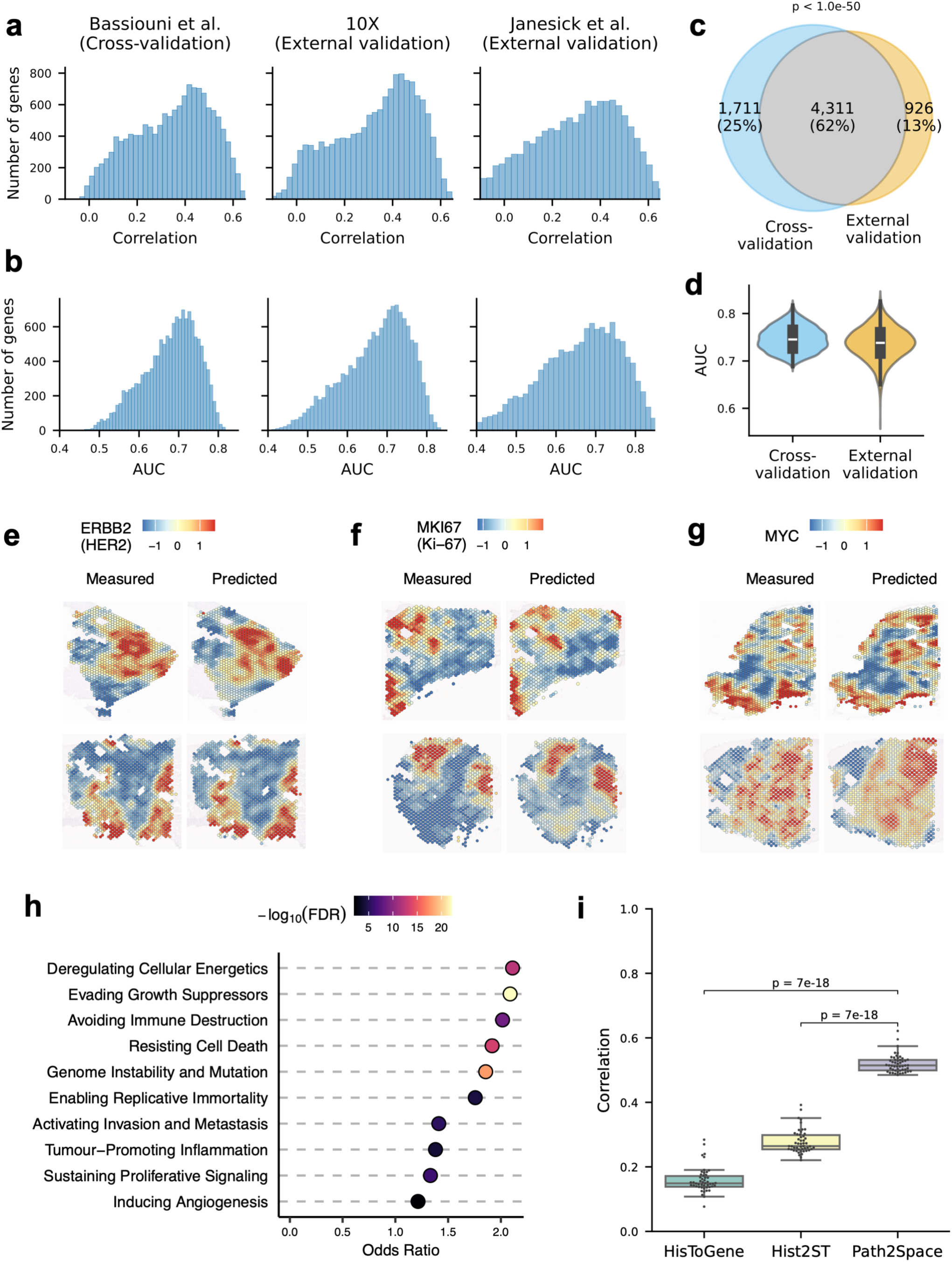
Performance of *Path2Space* in predicting spatial gene expression. **a,** Histograms of Pearson correlation coefficient (PCC) between predicted and actual gene expression values, showing cross-validation results from the Bassiouni et al. cohort (left) and independent validation in the 10x cohort (middle) and Janesick et al. (right) datasets. **b,** Histograms of AUC values for classifying high and low expression states across all genes. Results are shown for cross-validation in the Bassiouni et al. cohort (left), independent validation in the 10x cohort (middle), and the Janesick et al. dataset (right). **c,** Venn diagram showing genes with a predicted/measured PCC > 0.4 in both cross-validation and independent validation sets. PCCs were calculated as the mean gene-wise value across slides in the cross-validation set and across slides in both validation sets. **d,** Violin plot showing AUC values for classifying high and low expression states of the 4,311 genes robustly predicted in both the cross-validation and external validation cohorts. **e-g,** Examples of three robustly predicted and clinically important genes: *ERBB2* (HER2) (**e**), *MKI67* (Ki-67) (**f**), and *MYC* (**g**). Each panel shows two cross-validation set example slides with spots colored by scaled expression levels. The left column displays measured expression levels, and the right column shows *Path2Space*-predicted expression levels for slides in the same row. **h,** Gene set enrichment of the 10 key cancer hallmarks set in the 4,311 genes robustly predicted in all cohorts (shown in c). Each row represents a different cancer hallmark set, with the horizontal axis displaying the odds ratio of its enrichment. The color scale represents the -log10 adjusted enrichment *P* value; all gene sets were significant (adjusted *P* value ≤ 0.05). **i,** Boxplots comparing the performance of *Path2Space* with HisToGene and His2ST, evaluated using leave-one-out cross-validation on a HER2+ dataset (41 slides). The boxplots display the correlation coefficients between predicted and actual gene expression values for the top 50 genes per slide across each method (as those were reported in the original publications). *P* values from the two-sided Mann-Whitney U-test.

To further study and validate the performance of *Path2Space* and demonstrate its generalizability, we applied the Bassiouni et al.-trained model to predict spatial gene expression profiles from H&E slide images in two independent ST cohorts: one from 10x consisting of 4 slides from 3 patients, HER2+, hormone receptor-positive (HR+), and one unknown, and another from Janesick et al. consisting of one slide from one HER2+ patient. (**Supplementary Fig. 2d**). The median PCC between actual and predicted expression values, averaged across all spots per slide and across the external validation cohorts, was 0.35 (**Fig. 2a**), with 5,237 genes exceeding a correlation coefficient of 0.4 (**Fig. 2a; Supplementary Fig. 1b**). Furthermore, to evaluate performance in classifying genes into high or low expression states in these external validation cohorts, we employed the previously described median-based binarization approach. This analysis yielded a median AUC of 0.68, with 1,803 genes achieving an AUC greater than 0.75 (**Fig. 2b**). Collectively, the closely aligned prediction performance metrics between cohorts demonstrate that *Path2Space* effectively generalized from a TNBC training cohort to predict gene expression in HER2+ and HR+ tumors within independent validation cohorts.

Remarkably, we observed substantial concordance in highly predicted genes between the Bassiouni et al.-trained model and the two validation cohorts: 62% (4,311/6,850) of genes achieving a correlation >0.4 in one cohort also surpassed this threshold in the other (**Fig. 2c**). For these well predicted genes, the model also demonstrated strong classification performance, achieving median AUCs of 0.75 and 0.74 in the cross and external validation cohorts, respectively (**Fig. 2d**). This group included genes that are crucial in breast cancer prognosis and treatment decisions, such as *ERBB2* (HER2), *MKI67* (Ki-67), and *MYC* (**Fig. 2e-g**). Remarkably, Gene Set Enrichment Analysis (GSEA) of the 4,311 concordant genes (correlation >0.4 in all three cohorts) revealed significant enrichment (FDR<0.05) for all 10 of the 10 well known cancer hallmark processes. Notably, pathways related to avoiding immune destruction, evading growth suppressors, and genome instability and mutation showed the strongest enrichment (**Fig. 2h**).

Given that *Path2Space* was trained exclusively on fresh frozen (FF) slides from the ST Bassiouni et al. dataset and that the external validation datasets consist of both FF slides and formalin-fixed paraffin-embedded (FFPE) slides, we assessed its performance on FF slides and FFPE slides separately. Median gene expression correlations ranged from 0.32 to 0.40 for both slide types (**Supplementary Fig. 1c**), indicating that *Path2Space* maintained strong predictive performance on FFPE slides, comparable to the FF samples. This observation suggests *Path2Space* is well-suited for application to both fresh and archival tissue specimens.

Finally, for comparison, we benchmarked *Path2Space* against two state-of-the-art approaches for predicting spatial gene expression from tissue slides: HisToGene^33^ and Hist2ST^21^. To ensure a fair comparison, we trained a *Path2Space* model on the same legacy spatial transcriptomics dataset of 36 slides from 8 breast cancer patients^3^ used by these methods, employing an identical leave-one-out cross-validation strategy. We further evaluated the correlation between our predictions and the experimental measurements across each spot, consistent with the approach used by the other methods. We compared the models using the metric reported in these publications: the mean correlation of the top 50 predicted genes. *Path2Space* markedly outperformed both models, achieving a median correlation of 0.51, compared to 0.15 and 0.26 for HisToGene and Hist2ST, respectively (**Fig. 2i**). Importantly, we compared our predictions to the raw (unsmoothed) measured expression values per spot for PCC calculation to ensure fair comparison with benchmark methods. Additionally, we found that smoothing the inferred spatial gene expression patterns further improved prediction accuracy for *Path2Space* (**Supplementary Fig. 1d**). These findings underscore *Path2Space’s* capability to accurately infer spatial gene expression patterns from H&E images of breast cancer, indicating its potential for biomarker discovery in large cohorts.

### Inferring spatial cell type composition from the predicted spatial transcriptomics

Visium’s 55 μm spots typically capture signals from multiple cell types, making further cellular decomposition important for characterizing the tumor spatial landscape. As described in the overview, we first trained *Path2Space* to predict spatial gene expression at the spot level from H&E images. We then utilized localized predicted gene expression to estimate cell type fractions, leveraging cell type annotated ROIs images from the PanopTILs dataset.

We employed *Path2Space* to infer gene expression for each annotated region in the PanopTILs dataset. We then developed an MLP regressor using 5-fold cross-validation, grouped by patients, to estimate fractions of cancer cells, lymphocytes, and stromal cells. Model performance was evaluated using correlations between actual and predicted cell type fractions across all ROIs. Our model achieved PCC of 0.82, 0.74, and 0.58 for cancer cells, lymphocytes, and stromal cells, respectively, with the lower correlation for stromal cells potentially attributable to their lower abundance in the dataset (**Fig. 3a**). Notably, GSEA revealed significant enrichment (FDR<0.05) of breast cancer signature genes as features for cancer cell fraction prediction and lymphocyte signatures for lymphocyte prediction. While the stromal signature exhibited the highest enrichment score for stromal prediction, it did not reach statistical significance (**Fig. 3b; Methods**).

**Fig. 3:**
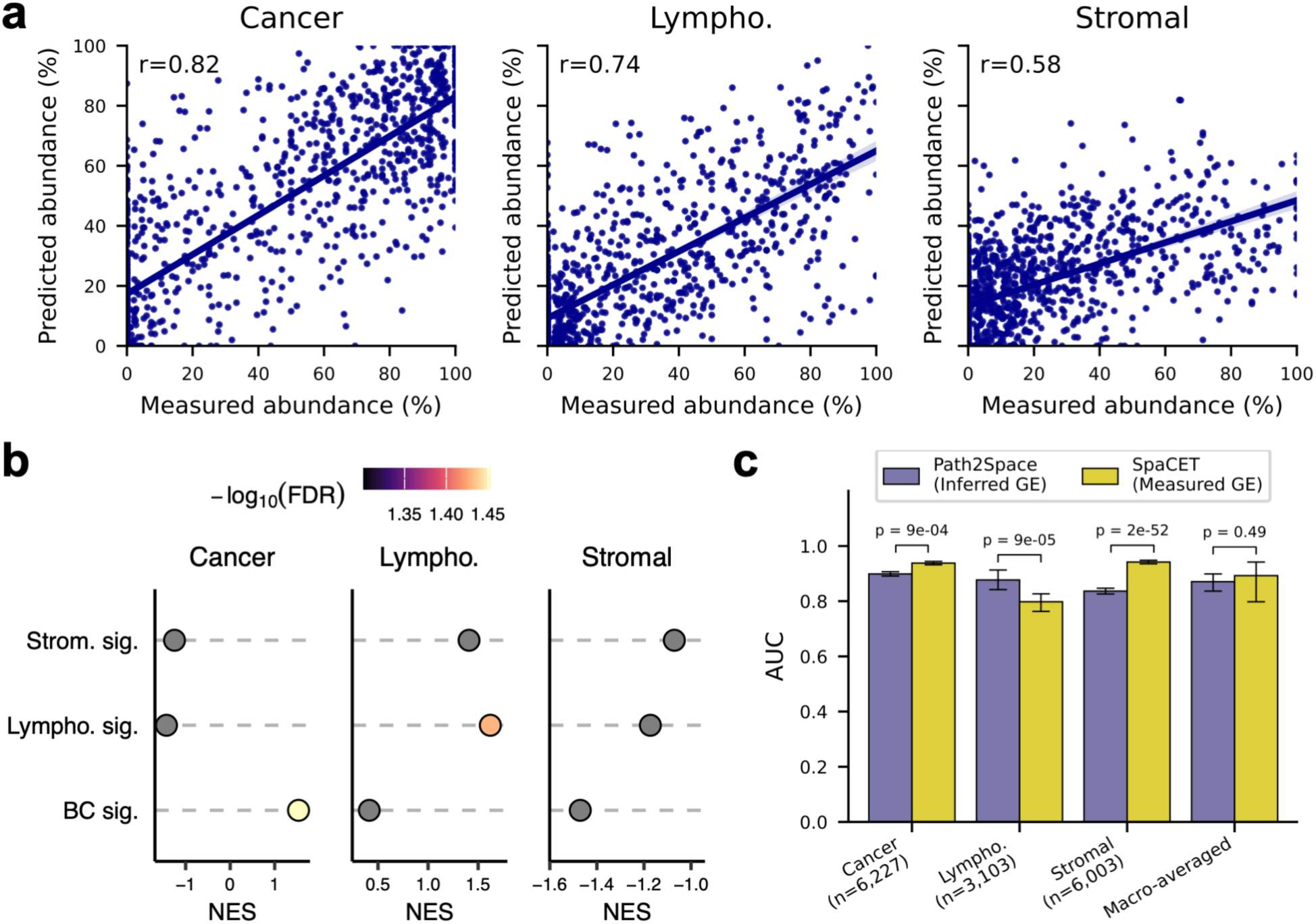
*Path2Space* performance in estimating cell type fractions from inferred spatial gene expression. **a,** Scatter plots showing cross-validation performance in the breast cancer patients PanopTIL dataset. Actual values are on the x-axes and predicted values on the y-axes for cancer cells (left), lymphocytes (Lympho., center), and stromal cells (right). PCC coefficients between actual and predicted values are displayed in each plot. **b**, GSEA showing cell type signature enrichment scores for each MLP prediction node, from left: cancer, lymphocyte, and stromal cells. (**Methods**). The x-axis shows the Normalized Enrichment Score (NES). The color scale indicates -log10 adjusted *P* value, with gray denoting non-significant values (FDR≥0.05). **c**, Performance comparison of deconvolution accuracy of *Path2Space* (from inferred gene expression) and SpaCET (using measured gene expression). The y-axis shows the area under the receiver operating characteristic curve (AUC) for cell type fractions predicted by *Path2Space* (purple) and SpaCET (orange), with micro-averaged AUCs shown rightmost. Error bars represent 95% confidence intervals derived from 1,000 bootstrapping iterations. Center points correspond to the computed AUC values for each model. *P* values were calculated using two-sided DeLong’s test for individual cell types and bootstrap testing (1,000 iterations) for macro-averaged AUCs.

To assess the performance of *Path2Space* for cell-type abundance estimation using external validation data, we applied the pipeline to two ST samples from the 10x dataset with spot-level pathologist annotations. One of the slides was derived from a HER2+ patient, while the other was from a patient with an unknown subtype (samples TENX13 and TENX39; **Supplementary Fig. 2e**). Using the PanopTILs-trained deconvolution model on the smoothed gene expression inferred from the slide images, we obtained AUC scores of 0.9, 0.87, and 0.84 for cancer cells, lymphocytes, and stromal cells, respectively. Further analysis revealed that smoothing the H&E-inferred spatial gene expression improved deconvolution accuracy (**Supplementary Fig. 3**). Remarkably, our H&E-inferred gene expression-based deconvolution model showed comparable performance to that of SpaCET^34^, which performed deconvolution using the *measured* gene expression as input. While SpaCET outperformed *Path2Space* for cancer and stromal cells, it demonstrated inferior performance for lymphocytes (**Fig. 3c**). *Path2Space* achieved a macro-averaged AUC of 0.87 across all cell types, comparable to the 0.89 achieved by SpaCET, with no significant difference between method*s* (*P* = 0.486, **Fig. 3c**). This comparable performance in cell identity decomposition between *Path2Space*’s H&E-based approach and methods based on measured ST underscores the potential of our algorithm to leverage widely available H&E images, thereby broadening the accessibility of spatial transcriptomics analysis.

### Breast cancer tumors are composed of three *spatially-grounded* subgroups with distinct TMEs and survival

Having established the ability of *Path2Space* to infer spatial gene expression and cell types from H&E-stained images, we applied the model to predict spatial gene expression from the slides in the TCGA breast cancer cohort, comprising 1,096 samples from 1,043 patients. The main goals of this analysis were two-fold: (1) From a basic science perspective, to characterize the TME of each sample and obtain an overview of breast cancer TME on an unprecedented scale. (2) From a translational angle, to identify *spatially oriented* prognostic biomarkers of breast cancer patient survival.

To emulate the spatial resolution of Visium training data, we divided each TCGA slide into contiguous "pseudo-spots" covering comparable areas, with an average of 6,131 pseudo-spots per slide, without the typical 45-90 micrometer gaps between Visium spots.

After inferring spatial gene expression for the entire cohort, we employed SpaGCN^35^, a graph convolutional network, to group pseudo-spots into continuous spatial domains based on their inferred transcriptomic profiles and spatial organization. This approach aims to capture continuous regions in the TME with similar as possible molecular, functional and cell type compositions. To achieve consistent cluster labeling across patients, the clustering was performed across all TCGA slides, identifying 11 shared spatial transcriptomic clusters (**Fig. 4a,b; Supplementary Fig. 4a-c**). These clusters, referred to as *ST clusters*, provide a unified framework for characterizing the TME in breast cancer. The ST clusters showed subtype-specific patterns (*P* = 0.003; **Supplementary Fig. 4d,e**), reflecting divergent TME organizations across breast cancer subtypes. Further analysis of these clusters revealed distinct patterns in the expression of breast cancer oncogenes and tumor suppressor genes, along with cluster specific activities of cancer hallmark pathways, cell-cell interactions and cell type compositions (**Fig. 4c,d; Supplementary Fig. 4f-h**). Notably, the abundance of certain ST clusters was found to be associated with disease-free survival (**Fig. 4e**).

**Fig. 4:**
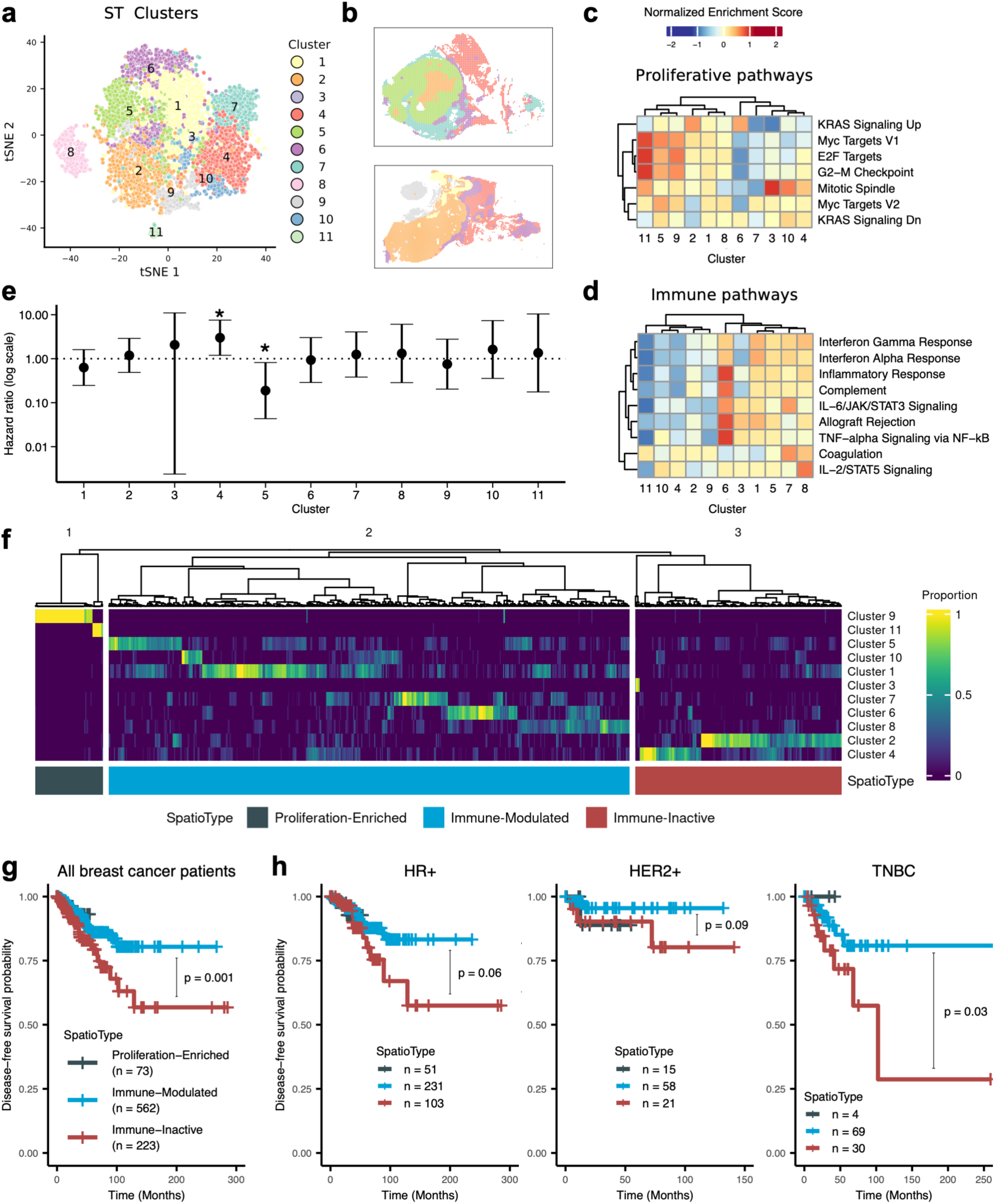
Identification of prognostic spatial tumor *SpatioTypes* in the TCGA BRCA cohort. **a,** t-SNE visualization of pseudo-spot gene expression profiles across TCGA slides, revealing 11 distinct spatial transcriptomic clusters. **b**, Two examples of TCGA slides visualizing their ST cluster assignments across the 11 shared clusters**. c,d,** Heatmaps showing NES of proliferative (**c**) and immune (**d**) pathways across ST clusters, based on MSigDB Cancer Hallmark gene sets. **e,** Cox regression analysis of individual spatial clusters reveals that the abundance of clusters 4 and 5 is significantly associated with survival (*P* < 0.05) across all patients. The vertical axis represents the hazard ratio on a log scale, with an asterisk (*) indicating statistical significance. **f,** Hierarchical clustering of spatial cluster abundances across *n* = 858 TCGA breast cancer samples reveals three distinct tumor *SpatioTypes*. The heatmap displays patients as rows and clusters as columns, with values representing spatial cluster proportions. The three spatial *SpatioTypes*— Proliferation-Enriched, Immune-Modulated, and Immune-Inactive—denoted by the color bar at the bottom. **g,** Kaplan-Meier survival analysis of disease-free survival across the three spatial *SpatioTypes* (*P* = 0.01, log-rank test). Patients in the Immune-Inactive *SpatioType* (n = 233) exhibit significantly poorer prognosis, particularly compared to the Immune-Modulated *SpatioType* (*P* = 0.001 shown in figure, single-sided log-rank test). **h,** Subgroup survival analysis shows that the poor prognosis of the Immune-Inactive *SpatioType* is consistent across the conventional HR+ (ER+ or PR+), HER2+, and TNBC subtypes, though subgroup sample sizes limit statistical significance. *SpatioTypes* colors correspond to panels (**f**) and (**g**), with the legend indicating patient counts for each *SpatioTypes*. *P* values from the single-sided log-rank test.

Based on the representation of each sample in terms of its cluster proportions, we performed hierarchical clustering of patient tumors. This analysis revealed three distinct spatially-grounded subgroups of breast cancer patients, termed *SpatioTypes*, each characterized by similar spatial cluster compositions (**Fig. 4f**). The sample-level clustering was optimized using the elbow method (**Supplementary Fig. 5a**). Among the three *SpatioTypes*, the TME of the **Proliferation-Enriched** *SpatioType* is characterized by ST clusters 9 and 11, which show strong activation of proliferative pathways (**Fig. 4c,f**). The **Immune-Modulated** *SpatioType*, the most prevalent in the TCGA cohort, features clusters with relatively high immune activity. In contrast, the **Immune-Inactive** *SpatioType* is dominated by clusters 2 and 4, marked by low immune activation (**Fig. 4d,f**).

In alignment with these spatial characteristics, TCGA measured bulk RNA-seq data reflected the molecular profiles of the *SpatioTypes*: Proliferation-Enriched patients showed strong activation of multiple proliferative hallmark pathways, while Immune-Modulated and Immune-Inactive patients generally showed high and low immune hallmark pathway activation, respectively (**Supplementary Fig. 5b**). Independently, analysis of TCGA TIL maps^36^ showed significantly higher proportions of tumor-infiltrating lymphocytes (TILs) in Immune-Modulated versus Immune-Inactive tumors, with mean TIL proportions 24% higher, further supporting their immune activation profile (*P* = 0.04, single-sided Mann-Whitney U-test; **Supplementary Fig. 5c**). Furthermore, the Proliferation-Enriched *SpatioType* was associated (*P* = 0.006, two-sided chi-squared test) with tumors of higher pathological stage (>1), consistent with its increased proliferative signature (**Supplementary Fig. 5d**).

Survival analysis of the 853 patients with both survival and slide data revealed that these three spatial *SpatioTypes* markedly stratify disease-free survival (*P* = 0.01, two-sided log-rank test). The comparison between the two most prevalent groups showed significant differences in survival between Immune-Modulated and Immune-Inactive patients (*P* = 0.001, two-sided log-rank test; **Fig. 4g, Supplementary Fig. 5e**). Notably, the Immune-Inactive *SpatioType* (n = 233) was indeed associated with poorer prognosis, even after adjusting for age and tumor stage, as one would expect given its ‘cold’ TME (**Supplementary Fig. 5f**). The Immune-Inactive *SpatioType* was mainly comprised of cluster 2, with high cancer cell density and oncogene upregulation, and cluster 4, a stromal cluster showing tumor suppressor downregulation - suggesting coordinated programs promoting tumor progression (**Supplementary Fig. 4f,g**). Remarkably, this spatially driven survival stratification complements and builds upon conventional breast cancer clinical subtype classifications, as the Immune-Inactive *SpatioType* was observed across HR+ (ER+ or PR+; n = 385), HER2+ (n = 94), and TNBC (n = 103) cohorts (271 patients lacked clinical annotations, **Methods**)^37^. However, statistical significance in subgroup analyses was limited by smaller sample sizes (**Fig. 4h, Supplementary Fig. 5g**).

### Inferred spatial gene expression patterns predict breast cancer patient survival and treatment response

Given that tumor heterogeneity is a key determinant of patient outcomes and therapeutic resistance^38^, we examine the prognostic value of *Path2Space*-inferred spatial gene expression patterns. To this end we employed a *SPAtial Neighborhood Diverseness (SPAND)* measure, which quantifies the mean variation of the expression of a gene or set of genes within a 100-µm^2^ neighborhood across the whole slide, relative to its overall mean expression. SPAND is computed by the negative Moran’s I^39,40^ of the gene scaled by the mean of the gene (**Methods**) across the slide and is the spatial analog of the classical standard deviation over the mean measure frequently used in statistics (**Fig. 5a**). High (low) SPAND scores denote high (low) heterogeneity. *Path2Space* inferred expression reliably captured this spatial heterogeneity measure (**Fig. 5b**), achieving a median correlation of 0.69 between SPAND values computed from predicted and measured ST for the 4,311 well-predicted genes across the 25 samples in the Bassiouni et al. cohort.

**Fig. 5:**
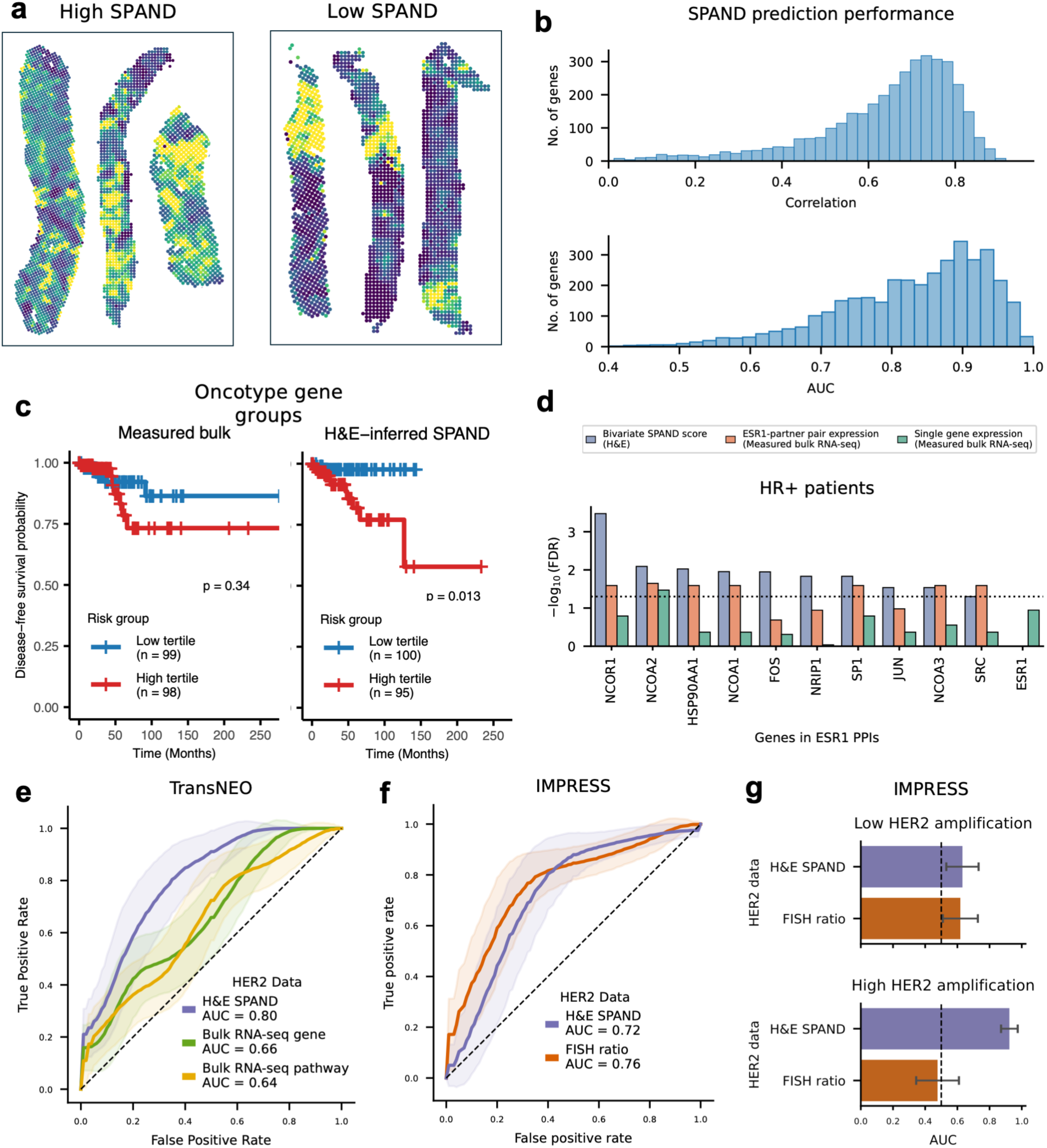
Spatial expression heterogeneity predicts survival and therapy response. **a,** Examples of TransNEO slides showing high (left) and low (right) HER2 SPAND scores. **b,** Distributions of the PCC (top, median = 0.69) and AUCs (bottom, median = 0.85) between the SPAND calculated based on measured and *Path2Space*-inferred ST data for all well predicted genes in the Bassiouni et al. cohort. **c,** Kaplan-Meier survival curves comparing the predictive power of the *measured* bulk RNA-seq values (left) and the inferred SPAND scores (right) for the OncotypeDX (n = 295) signatures in the TCGA. Risk groups show top and bottom tertiles; *P* values from log-rank test. **d,** Association with overall survival of *ESR1* and its protein-protein interaction partners in TCGA HR+ breast cancer patients (n=385), comparing bivariate SPAND scores with bulk RNA-seq measurements of individual genes and mean expression of gene-*ESR1* pairs. **e-g,** ROC curves comparing HER2 SPAND scores versus *ERBB2* expression levels and HER2 pathway enrichment scores, the latter measured using the actual bulk RNA-seq in TransNEO (**e**). A similar comparison versus FISH measured HER2/CEP17 ratio in the IMPRESS cohort (**f**). The shaded areas depict bootstrapped-based standard errors (n=1,000); Area under the curve (AUC) values are provided in the legend. **g**, Response prediction AUCs comparing HER2 SPAND and HER2/CEP17 FISH ratios in IMPRESS cohort, stratified by HER2 amplification level. Patients were divided into low (HER2/CEP17 ≤ 6.73; upper panel) and high (HER2/CEP17 > 6.73; lower panel) amplification groups, using the cohort’s median HER2/CEP17 ratio of 6.73 as the threshold. Error bars: standard errors from 1,000 bootstrap iterations. Bar height shows observed AUC values.

Having established that, we next evaluated the potential of SPAND scores to enhance the predictive signal of extant clinically useful prognostic signatures, focusing first on the gene groups defined in the FDA-approved Oncotype DX^41,42^, a widely adopted and validated diagnostic assay. The Oncotype DX signature calculates risk based on the expression of 21 genes, grouped into functional categories, with coefficients optimized for measurements obtained using its proprietary reverse-transcriptase polymerase chain reaction (RT-PCR) assay. However, when applied to TCGA bulk RNA-seq data—which differs significantly from RT-PCR in measurement distributions—the original signature failed to stratify disease-free survival outcomes on the subset of patients who met the FDA clinical guidelines for Oncotype DX (**Methods, Supplementary Fig. 6a**). Furthermore, even after optimizing the coefficients to fit this signature to RNA-seq data (**Methods**), it was not predictive (C-index: 0.51; **Supplementary Fig. 6b**) and did not significantly separate disease-free survival between high- and low-risk tertiles (**Fig. 5c**). In contrast, SPAND-based models trained on the same TCGA cohort achieved significantly better performance, with higher prognostic accuracy (C-index: 0.60; **Supplementary Fig. 6b**) and significantly stratified disease-free survival between the top and bottom risk tertiles (**Fig. 5c**). Similarly, we extended this analysis to the FDA-approved MammaPrint^43,44^ signature (**Supplementary Fig. 6a)**. Once again, SPAND-based models outperformed bulk RNA-seq, achieving higher C-index values (**Supplementary Fig. 6b**) and better stratification of disease-free survival (**Supplementary Fig. 6c,d**). These findings underscore the potential of incorporating spatial information via SPAND scores to enhance the predictive power of extant prognostic signatures.

Despite the dependency of HR+ breast cancers on Estrogen Receptor (ER) activity, its measured bulk RNA-seq expression levels failed to show prognostic value in HR+ patients in the TCGA data (**Fig. 5d**). We therefore examined the prognostic value of spatial co-variation between these receptors and their direct protein interaction partners. To quantify the spatial co-variation, we introduced the *bivariate SPAND* (−Bivariate Moran’s Index scaled by mean expression)^45^. Analysis of ER gene *ESR1* and its protein-protein interaction (PPI) partners^46^ in the TCGA breast cancer cohort revealed that in HR+ (n=385), high bivariate SPAND scores of *ESR1* with each of its PPI partners were significantly associated with worse overall survival (FDR<0.05), outperforming bulk RNA-seq mean expression measurements (**Fig. 5e; Supplementary Fig. 7a,b**). Reassuringly, this association was not observed for TNBC or HER2+ breast cancer patients (**Supplementary Fig. 7a,c**).

Next, we investigated the ability of SPAND scores of HER2 to predict trastuzumab response. Given that HER2-high cells are more sensitive to trastuzumab-mediated immune response ^47–52^, we hypothesized that tumors with high SPAND, where HER2-high cells intermix with HER2-low cells, would show better response, as immune response elicited by targeted and dying HER2-high cells could influence nearby HER2-lower cells. To examine this hypothesis, we applied *Path2Space* to two breast cancer cohorts treated with trastuzumab, TransNEO (HER2+ n=61; HER2-n=93) and IMPRESS (HER2+ n=62; HER2-n=64). We inferred the spatial gene expression and cell type fractions in each slide to compute the SPAND scores of HER2 pathway activity normalized by cancer cell fraction (HER2 per cancer).

We found that HER2 pathway activity SPAND scores significantly predict complete pathological response (pCR) to trastuzumab in both TransNEO (AUC=0.8, *P* = 0.006, two-sided Mann-Whitney U-test) and IMPRESS (AUC=0.72, *P* = 0.004) cohorts, outperforming the mean inferred HER2 per cancer per tumor (**Supplementary Fig. 8a**). Moreover, the predictive power of HER2 SPAND is specific to trastuzumab, showing weaker association with chemotherapy response (AUC=0.61, *P* = 0.06 and AUC=0.57, *P* = 0.15; **Supplementary Fig. 8b**). Notably, HER2 SPAND predictive power outperformed bulk RNA-seq measurements in TransNEO (**Fig. 5e**). In IMPRESS, while showing comparable overall performance to HER2/CEP17 ratios measured by FISH (**Fig. 5f**), HER2 SPAND better discriminates pCR in cases of high HER2-amplification (AUC=0.92, *P* = 0.001; **Fig. 5g**).

Notably, previous studies used *non-spatial* metrics of HER2 heterogeneity vs homogeneity, finding that heterogeneity is associated with poor response to trastuzumab^53,54^. In contrast, SPAND captures distinct information by measuring variance/heterogeneity localized to neighborhoods of micrometer scale. Overall, these results highlight the added value of spatial information in defining heterogeneity on a microscopic scale. These results underscore the considerable potential added value of using spatially resolved gene expression analysis for identification of more accurate treatment-specific biomarkers.

### *Spatially-grounded clustering* further predicts patient treatment response to trastuzumab and chemotherapy

To further identify spatial biomarkers for treatment response, we developed and studied a complementary approach by leveraging the 11 breast cancer ST clusters identified earlier in our TCGA analysis. Following the exact same procedure, we used for inferring patient survival in the TCGA cohort, we aligned the 11 clusters with the pertaining TransNEO and IMPRESS cohorts to quantify the fraction (proportion) of each of the 11 clusters in each patient sample.

To predict chemotherapy response based on this 11-cluster representation of each tumor sample, we trained a logistic regression classifier using breast cancer ST cluster proportions as features in the TransNEO cohort. Five-fold cross-validation on chemotherapy-treated TransNEO patients (93 HER2-patients) achieved AUC of 0.75 and AUPRC of 0.45 (the overall response rate in this cohort was 0.23; **Fig. 6a; Supplementary Fig. 9c**). For comparison, we evaluated two alternative approaches using five-fold cross-validation on the TransNEO cohort: a direct image-based model that applied similar preprocessing and feature extraction followed by MLP classification, and an imputed bulk model that predicts transcriptomic profiles from H&E images, trained on BRCA TCGA RNA-seq data (**Supplementary Fig. 9a,b**), followed by logistic regression on the imputed gene expression to classify response. The direct (image-based) model showed lower performance (AUC = 0.69, AUPRC = 0.38), while the imputed bulk approach achieved a quite similar performance on this cohort (AUC = 0.73, AUPRC = 0.51; **Fig. 6a**). However, testing these three prediction models on another independent dataset, the IMPRESS cohort, consisting of chemotherapy-treated TNBC patients (n=64), their performance diverged. The *Path2Space* model achieved AUC of 0.74 and AUPRC of 0.74 despite being trained on a different mixed ER+ and ER-patients cohort, while the imputed bulk model failed to generalize (AUC=0.52; AUPR=0.44), as did the direct image-based model (AUC=0.57; AUPR=0.51). Moreover, the cluster-based *Path2Space* model outperformed five published H&E-based predictors evaluated on the IMPRESS cohort^55^, including the model by Huang et al.^32^ (AUC = 0.68), despite the latter being both trained and tested on IMPRESS through cross-validation (**Fig. 6a; Supplementary Fig. 9c**).

**Fig. 6:**
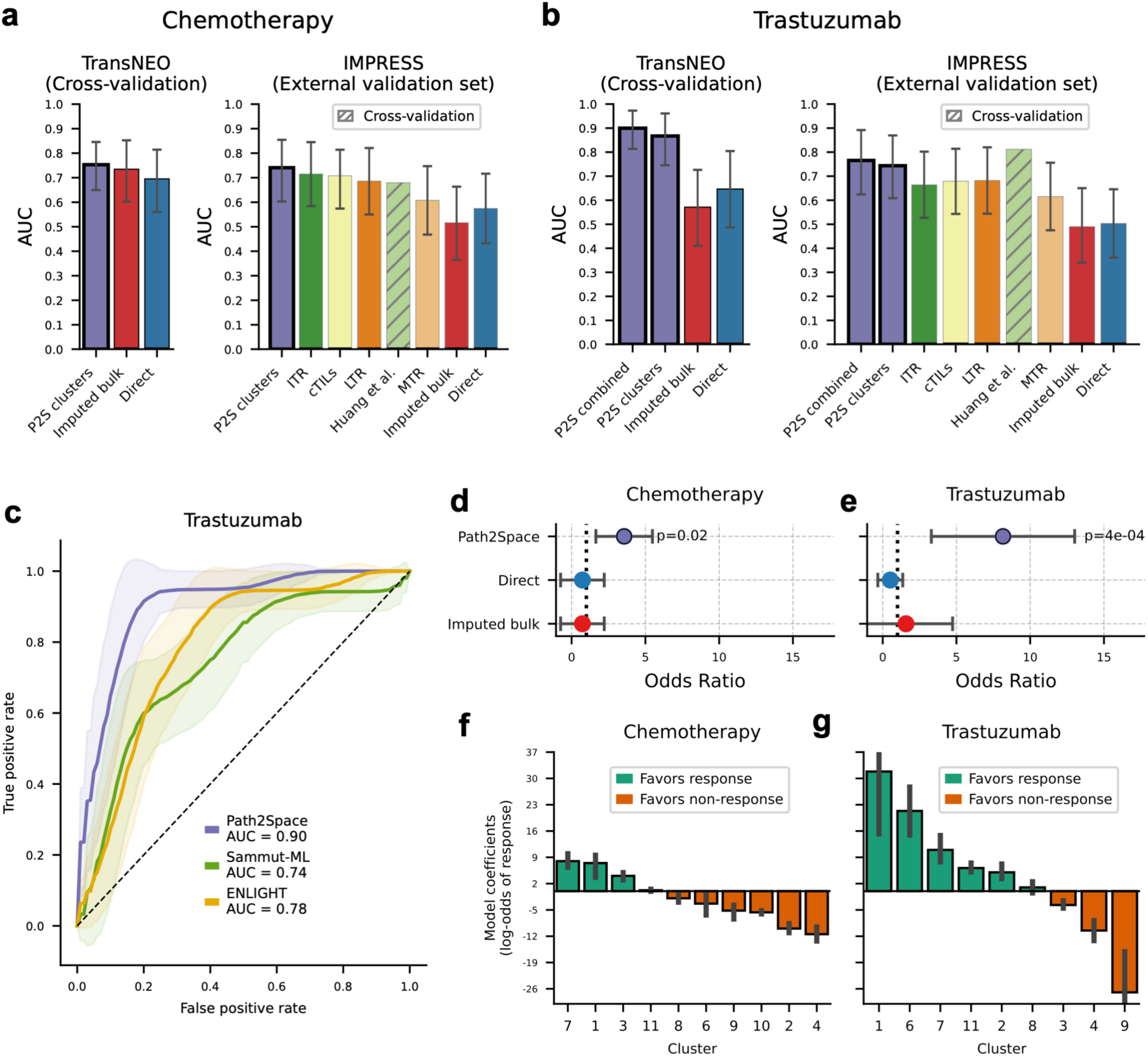
*Path2Space* predicts chemotherapy and trastuzumab response based on TME composition. **a,b,** AUC for response prediction in TransNEO (cross-validation, left) and IMPRESS (external validation, right) cohorts for chemotherapy (**a**) and trastuzumab (**b**), comparing *Path2Space* with other H&E-based models. *Path2Space* models (P2S, displayed with a thick bar edge) include a cluster model for both treatments and a combined model (integrating spatial clusters with HER2 per cancer SPAND) for trastuzumab. For IMPRESS we additionally show Huang et al.’s cross-validation performance and Aswolinskiy et al.’s H&E-based models: lymphocyte-tumor ratio (LTR), computational tumor infiltrating lymphocytes score (cTILs), inflamed tumor ratio (ITR) (orange), and mitoses-tumor ratio (MTR). Error bars represent the 95% confidence intervals computed over 1,000 bootstrap iterations. Bar height displays the resulting AUC values. **c,** ROC curves comparing *Path2Space* to models using the *measured* bulk omics data: Sammut-ML (incorporating RNA-seq, genomics, and clinical data) and ENLIGHT (RNA-seq) in the TransNEO trastuzumab cohort. Shaded areas denote the bootstrap standard errors (n=1,000); AUC values are provided in the legend. **d,e,** Odds ratios for response prediction in IMPRESS validation cohort: chemotherapy (**d**) and trastuzumab with combined model (**e**). * indicates *P* < 0.05 (Fisher’s exact test). **f,g,** Clusters contributions in logistic regression models for chemotherapy (**e**) and trastuzumab (**f**) response prediction. Mean coefficients from five-fold cross-validation are shown with standard errors; positive (green) and negative (orange) values indicate association with response and non-response, respectively.

We next turned to develop an ST-based predictor of response to trastuzumab, based on the 11-cluster representation. We trained a logistic regression model using breast cancer ST cluster composition from TransNEO’s trastuzumab-treated patients (n=61), with the trastuzumab-treated IMPRESS patients for independent validation. The model achieved AUC of 0.86 and AUPRC of 0.70 (overall response rate: 0.61) on TransNEO and AUC of 0.74 and AUPRC of 0.75 (overall response rate: 0.31) on IMPRESS, outperforming models based solely on image features or H&E-imputed bulk gene expression in both cohorts (**Fig. 6b, Supplementary Fig. 9d**). In the IMPRESS cohort (n=62), our model outperformed four of the same five H&E-based predictors, with only the model from Huang et al. showing slightly higher performance (but importantly, only through cross-validation) on the IMPRESS cohort (AUC=0.81). Finally, building on these results, the integration of our cluster-based model with the HER2 SPAND biomarker presented in the previous section further improved prediction accuracy, achieving an AUC of 0.90 and AUPRC of 0.78 on TransNEO and AUC of 0.77 and AUPRC of 0.8 on IMPRESS (**Fig. 6b, Supplementary Fig. 9d**).

We additionally compared our trastuzumab response models to recently reported models evaluated on TransNEO that use measured, non-spatial molecular data. Remarkably, the *Path2Space* models outperformed both Sammut-ML^31^ (AUC=0.74), which uses clinical, genomic, and bulk RNA sequencing data, and ENLIGHT^56^ (AUC=0.78), which relies on RNA-seq data (**Fig. 6c**). The integrated model’s improvement over Sammut-ML was statistically significant (*P* = 0.045, single-sided DeLong’s test). However, Sammut-ML showed superior performance in chemotherapy response prediction (**Supplementary Fig. 9e**). To study the potential clinical utility of the ST predictor, we used the TransNEO cohort to define decision thresholds using the Youden method^57^. We applied the resulting threshold to the IMPRESS cohort and found that patients with scores above the thresholds showed significantly higher odds of response, with odds ratios of 3.6 (*P* = 0.02) for chemotherapy and 8.2 (*P* = 4e-4) for trastuzumab. (**Fig. 6d,e**).

To derive further insights into the spatial determinants of treatment response, we analyzed the logistic regression coefficients of our trained models. In the chemotherapy model, cluster 4 showed the strongest association with poor response, followed by cluster 2 (**Fig. 6f**). These findings align with findings from our TCGA analysis, where both clusters were defining features of the poor-prognosis Immune-Inactive *SpatioType*, with cluster 4 exhibiting the strongest univariate association with poor disease-free survival (**Fig. 4e,f**). In the trastuzumab model, cluster 4 remained associated with poor response, albeit less strongly (**Fig. 6g**). Conversely, clusters 1 and 6, which are most strongly associated with better response, displayed above-average HER2 SPAND levels, with cluster 6 having the second-highest levels (**Supplementary Fig. 9f**).

## Discussion

In this study, we introduce *Path2Space*, a deep learning tool that robustly predicts the spatial gene expression of thousands of genes directly from histopathology slide images on an unprecedented level. Applied to the TCGA breast cancer cohort, it identified three breast cancer subgroups defined by spatial transcriptomic patterns - termed *SpatioTypes* - that significantly stratify disease-free survival. Applying *Path2Space* to two chemotherapy and trastuzumab response cohorts revealed spatial markers significantly associated with treatment outcomes, showcasing its potential for identifying clinically relevant biomarkers and enhancing our ability to predict patient response speedily in an equitable manner directly from patient pathology slides.

Our investigation into trastuzumab response employed two complementary approaches, each providing valuable insights. The first is based on measuring the spatial SPAND scores of established response biomarkers. Here we found that tumors with high SPAND scores of HER2 expression, designating mixed HER2-high and HER2-low spatial regions, would be more likely to respond to treatment. This finding is in concordance with previous reports about trastuzumab’s ability to mobilize immune effector cell cytotoxicity through Fc receptor engagement^47^, which may in turn influence the surrounding microenvironment. These findings complement previous studies on HER2 heterogeneity^53,54^, providing further testimony that (inferred) HER2 spatial heterogeneity is indeed predictive of response.

Our second approach is based on building representations of individual tumors by the proportions of spatial clusters composing them. Our analysis of the TCGA breast cancer cohort identified and characterized 11 spatial transcriptomic clusters. This cluster based representation enabled the development of simple machine-learning models to predict treatment response directly from H&E slides. Examination of these models highlighted specific spatial environments strongly associated with clinical outcomes, offering valuable insights into their underlying molecular characteristics and cell-cell interactions.

While *Path2Space* shows great promise in predicting spatial gene expression from histopathology slide images, it is not without limitations. Our findings indicate that the correlation between measured ST and predicted gene expression is high for only a subset of approximately 4,300 highly expressed genes. This puts bounds on the ability of our model, at least in its current form, to capture the full spectrum of gene expression within the tumor and its microenvironment. Additionally, the cell type fraction estimation analysis based on *Path2Space* inferred ST is still slightly less accurate compared to deconvolution based on measured ST data, as expected. This discrepancy highlights the continuing importance of conducting spatial transcriptomics experiments for precise cellular mapping. Nevertheless, bearing those limitations in mind, our results clearly demonstrate that the level of accuracy achieved by *Path2Space* is sufficient for inferring spatial gene expression and cellular composition from whole slide images (WSI) at a high-resolution tile level, enabling the analysis of much larger datasets than is currently feasible with ST methods.

As the amount of spatial transcriptomics data continues to grow, with higher read counts and finer spatial resolution—even at a single-cell level^58^—we expect improved predictions of spatial gene expression and more precise inferences of cellular composition. Notably, while trastuzumab and chemotherapy response served as a test case, the *Path2Space* approach is broadly applicable. It can be extended to any treatment where H&E images and response data are available, offering the potential to revolutionize biomarker discovery and treatment response prediction across diverse cancer types and therapies. Furthermore, the model’s framework is adaptable to other cancer types, offering the potential for broader insights into different TMEs. Beyond transcriptomics, *Path2Space* could be readily expanded to integrate other kinds of spatial omics data, such as methylation and proteomics^59^, as these measurement technologies become available.

Ultimately, *Path2Space* provides a scalable, rapid and cost-effective solution for predicting spatial gene expression, facilitating the analysis of large datasets previously inaccessible to spatial transcriptomics methods. By unlocking the potential of spatial transcriptomics at scale, it can generate unprecedented insights into tumor architecture and its molecular foundations, paving the way for more personalized and effective treatment strategies. By facilitating the analysis of larger patient datasets, *Path2Space* extends the reach of spatial transcriptomics to advance precision oncology, with the potential to significantly improve patient outcomes.

## Methods

### Data collection

All datasets used in this study are from publicly available sources. Links to download the datasets can be found in the Data Availability section below.

### Spatial transcriptomics data preprocessing

For model training, gene expression values were incremented by 1 and log10-transformed. Due to the high proportion of missing values typical in spatial transcriptomics data, we focused on 14,068 genes that were detected in at least 5% of spots on each slide in the Bassiouni et al. dataset, ensuring sufficient data for robust training across genes.

Given our smoothing procedure (see below), we restricted training to slides where, on average, each spot had eight neighboring spots within a 200-micrometer distance, the minimum spacing between spots. This filtering resulted in a final set of 25 slides from the Bassiouni et al. dataset used for model training.

### *Path2Space* computational framework

#### Spatial gene expression prediction

The architecture consists of three main components (**Fig. 1a**):

1. **Image Processing:** The preprocessing phase began by dividing whole-slide images into non-overlapping 224×224 pixel tiles centered around each spatial transcriptomics spot. Tiles with more than 50% background content are excluded (QC). This process resulted in several thousand tiles per slide, depending on the slide’s dimensions. To address staining variability, color normalization techniques^16^ were applied.
2. **Feature extraction:** Tile images corresponding to spots were processed using CTransPath^27^, a foundational digital pathology model trained through self-supervision on histopathology images. Each tile was represented by a 768-dimensional CTransPath feature vector.
3. **Multilayer Perceptron (MLP) Regression:** This component built a predictive model linking CTransPath features to whole-genome spot-level gene expression. The model architecture includes three layers: (1) an input layer with 768 nodes, corresponding to the size of the CTransPath feature vector; (2) a hidden layer with 768 nodes; and (3) an output layer with 14,068 nodes, each representing a gene. ReLU activation is applied to each layer to introduce non-linearity and enhance model performance.

#### Spatial smoothing

Gene expression values were smoothed by averaging each spot’s expression with that of all its neighboring spots within a 200-micrometer radius, the minimum distance between spots in the dataset. On average, this resulted in 8 neighbors per spot being used for smoothing. This procedure was applied to both model predictions and original expression measurements (**Supplementary Fig. 1a**) and was used for model evaluation and downstream analyses.

#### Model training procedure

##### Spatial gene expression prediction

The models were trained using stochastic gradient descent with mini-batch processing. For optimization, we employed the Adam optimizer with an initial learning rate of 0.0001. The mean squared error (MSE) between predicted and actual values was used as the loss function.

Training of the MLP regressors and classifiers was conducted over 500 epochs. To reduce the risk of overfitting, a dropout rate of 20% was applied to the initial layer. We also implemented early stopping to enhance computational efficiency and further prevent overfitting. The training was stopped if no improvement was observed in the mean correlation between predicted and actual gene expression values on the validation set after 50 epochs.

#### Cross-validation

Our training and evaluation strategy followed the protocol established in our previous work^16,17^, employing a leave-one-patient-out cross-validation approach. We implemented a 15×7 nested cross-validation scheme, where each of the 15 outer folds used slides from one patient for testing. Within each outer fold, seven inner folds were created, with slides from a single patient designated for validation.

For generating cross-validation predictions on the Bassiouni et al. training dataset, we averaged the outputs from the seven models, each trained on a different inner fold, for the patient slides in the respective outer fold. For the external validation datasets and clinical H&E cohorts, predictions from all 15×7=105 models were averaged to produce the final prediction.

#### Model evaluation

We evaluated gene-wise prediction accuracy by first calculating the Pearson correlation between spatially smoothed actual and predicted expression values across all spots within each slide. We then averaged these correlations across genes and slides from the same patient. Finally, for each gene, the correlation values were averaged across all patients. PCC were computed using SciPy^50^.

We also assessed our ability to classify high or low gene expression. We binarized the gene expression labels according to the median value per slide, and then calculated the AUCs for the prediction of the binarized gene expression for each gene, using the predicted gene expression values as classification scores. Similarly to PCC calculation, for a given gene, we first averaged AUCs across slides from the same patient, and then averaged across all patients.

#### Comparison to HisToGene and Hist2ST

To ensure rigorous benchmarking, we retrained our model using the same legacy spatial transcriptomics dataset of 36 slides derived from 8 HER2+ breast cancer patients, as employed by HisToGene and Hist2ST. We adhered to the cross-validation approach described in these studies, utilizing a leave-one-slide-out cross-validation strategy. This involved 36 outer folds, corresponding to each slide, and 7 inner folds for model tuning.

For performance comparison, we computed the PCC for the top 50 genes, replicating the evaluation metrics reported in the publications of HisToGene and Hist2ST. The gene list and PCC values for these benchmarks were sourced directly from the respective publications.

### Spatially resolved estimation of cell type fractions

To estimate cell type fractions at Visium spot resolution, we leveraged the PanopTILs dataset, which includes individual nuclei annotations from 1,709 regions of interest (ROIs) across 151 TCGA breast cancer patients. This dataset provided cell type identities for three prevalent cell types: cancer cells, lymphocytes, and stromal cells.

#### Pseudospot extraction and model training

Tile images matching the size of Visium spots (pseudospots) were extracted from each ROI, and the fractions of each cell type within each image were computed based on their nuclei annotations. To train and evaluate the predictor, we used *Path2Space* to generate spatial gene expression profiles for the pseudospot images. These gene expression profiles served as input to a MLP regressor tasked with predicting cell type fractions.

#### Model development and optimization

Within each training fold, we normalized *Path2Space*-inferred gene expression data to a range of 0–1 using Scikit-learn’s MinMaxScaler. We then selected the top 1,000 genes with the highest variance across tumor classes, applying ANOVA F-statistics with SelectKBest and f_classif.

The MLP regressor was designed with three output nodes, predicting the fractions of cancer cells, lymphocytes, and stromal cells. Hyperparameters—including learning rate, backpropagation algorithm, momentum, activation functions, number of layers, iterations, and validation split— were fine-tuned through five-fold cross-validation within the training fold. This optimization process, guided by the regression F-score, was performed using Scikit-learn’s GridSearchCV.

#### Cross-validation

We used 5-fold cross-validation, keeping all slides from each patient in the same fold to prevent data leakage and ensure model generalizability. For TCGA cross-validation assessment, we combined predictions from all test folds and computed PCC. For test set predictions, we averaged the outputs from all five models to estimate cell type fractions.

#### Performance evaluation

Predicted cell type fractions and labels from each fold were aggregated across all test folds to evaluate performance. For external validation, predictions were averaged from the five cross-validation models and applied to H&E images and cell type annotations from the 10x cohort (**Fig. 1b**).

#### Model Interpretation and Feature Attribution

To analyze model performance and identify key predictive genes, we aggregated feature sets (genes) from all five trained models (across each of the five folds). Feature importance for each cell type was computed using Feature Attribution via Weight Propagation—a method of linear model interpretation through layer-wise weight analysis ^60^. For genes appearing in more than one fold, importance scores were averaged across models.

#### Pathway enrichment analysis

We tested gene enrichment using cell type signatures from the SpaCET^34^ R package, focusing on breast cancer, lymphocytes, and stromal cells (CAFs and macrophages). From each signature, we selected the top 50 most highly expressed genes. Using GSEAPY^61^, we performed pre-ranked GSEA by ranking genes based on their activation scores and using SpaCET cell type signatures as gene sets. We then assessed enrichment by testing the activation scores from each cell type against all three signatures.

### Analysis of predicted spatial gene expression

For downstream analysis of the predicted spatial gene expression, we utilized smoothed values for 4,311 genes that exhibited strong correlations (above 0.4) in both the Bassiouni et al. and 10x datasets.

Gene set enrichment analysis was performed using the Python package GSEAPY, with the prerank function^62^ for NES and the enrichr function for overrepresentation analysis (**Fig. 2h**).

### Spatial transcriptomic (ST) clusters

For clustering, we first applied SpaGCN^35^ to the inferred spatial gene expression data to identify slide-specific spatial domains — regions within each slide that are spatially and gene expression coherent. To ensure consistent cluster labeling across tumor slides from different TCGA patients, we averaged the gene expression of each domain across all TCGA slides. The data were normalized using Seurat’s log-normalization method.

Next, we employed Seurat’s^63^ *FindClusters* function to group these domain-averaged expression profiles, identifying 11 spatial clusters that were consistent across the TCGA cohort. Each domain was assigned to one of these 11 transcriptomic clusters, resulting in unified cluster labels that were spatially and biologically consistent across all samples (**Supplementary Fig. 4a-c**).

To transfer these spatial clusters to independent datasets (TransNEO and IMPRESS), we followed the same pipeline. First, we applied SpaGCN to infer spatial domains from gene expression data in the TransNEO and IMPRESS cohorts, which contained treatment response data for both chemotherapy and trastuzumab. To align domain expression profiles between each cohort and TCGA, we used Seurat’s anchor-based integration method (TransferData function), mitigating batch effects and ensuring consistent spatial cluster assignment.

#### Differential expression analysis

To assess differential gene expression across ST clusters, we calculated the relative expression of each gene within a cluster compared to the mean expression across all other clusters within the same slide. For each slide, the mean expression of each gene was computed per cluster. The fold change was then determined by comparing the cluster-specific mean to the overall slide mean. Linear regression was used to assess statistical significance and estimate fold change (R version 4.3.2, stats package version 4.3.2).

#### Pathway enrichment analysis

A similar approach was applied for pathway enrichment analysis. Instead of raw gene expression values, we utilized single-sample gene set enrichment analysis (ssGSEA), implemented in GSEAPY, to transform the expression profiles into NES based on the MSigDB Cancer Hallmark gene sets. Pathway scores were averaged per cluster and slide, and fold changes were computed relative to the slide’s overall pathway activity using linear regression.

#### Cell type composition

To determine the cell type composition of ST clusters, predicted cell type abundances were averaged across all slides within each cluster. The resulting cluster-level cell type fractions were further averaged across clusters to obtain a global cell composition profile for each cluster.

#### Ligand-receptor interaction analysis

We applied a ligand-receptor interaction analysis inspired by the differential analysis pipeline from SpatialDM^64^, utilizing a z-score-based method. Global z-scores of bivariate Moran’s I statistics were computed for each slide using SpatialDM, quantifying the spatial co-expression of ligand-receptor pairs across tumor regions.

For each ligand-receptor interaction, a linear regression model was fit, with cluster proportions serving as the independent variables (features) and the interaction z-scores as the response. Significant ligand-receptor pairs were identified through chi-square goodness-of-fit tests, applying a false discovery rate (FDR) threshold of < 0.05. The regression coefficients were further used to estimate the extent of ligand-receptor co-expression within individual clusters, offering insights into spatial interaction dynamics throughout the tumor microenvironment.

#### Hierarchical clustering for identifying *SpatioTypes*

We performed hierarchical clustering on the patient-cluster proportion matrix to identify distinct *SpatioTypes*. First, we standardized the input matrix using z-score scaling for each feature (cluster) to ensure equal contribution to the clustering process.

We computed a distance matrix using Euclidean distances between the scaled feature vectors. Agglomerative hierarchical clustering was then performed using Ward’s minimum variance method (ward.D2), which minimizes the total within-cluster variance at each merging step to produce compact, spherical clusters.

To determine the optimal number of clusters (k), we calculated the within-cluster sum of squares (WSS) for k ranging from 1 to 10. For each k, we cut the hierarchical clustering dendrogram to form k clusters and computed the WSS by summing the squared differences between data points and their respective cluster centroids. We used the elbow method to visualize the relationship between k and WSS, identifying k=3 as the optimal k at the point where the WSS reduction rate substantially decreased.

All analyses were performed in R (version 4.3.2) using the factoextra (version 1.0.7) and stats packages.

#### Survival analysis and clinical annotations

We analyzed overall and disease-free survival data from TCGA^65^. Survival analyses were performed using the survival package (version 3.5.7) in R, implementing log-rank tests and Cox proportional hazards models.

For clinical subtype annotations, we used receptor status data from TCGA^37^. We classified patients as HER2+ when HER2 status was marked ’positive’. Among HER2-patients, those with positive status for either ER or PR were classified as HR+. Patients negative for all three receptors (HER2, ER, and PR) were classified as triple-negative breast cancer (TNBC). We excluded patients with missing receptor status data or those marked as ’equivocal’ in TCGA annotations.

### SPAND analysis

#### SPAND calculation

SPAND was defined as 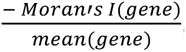, where the Moran’s I and the mean were calculated on the predicted expression of a given gene across pseudospots. Moran’s I, which quantifies spatial autocorrelation, was computed using the PySAL^66^ (24.01) package (esda module) with spatial weights derived from the pseudospot grid. The weights matrix was generated with the lat2W function, applying a rook’s case contiguity structure based on the grid dimensions (number of rows and columns), connecting each pseudospot to its four cardinal neighbors (top, bottom, left, right). To ensure robustness and comparability across spots with varying numbers of neighbors, the weights matrix was row-standardized, ensuring equal contributions to the final Moran’s I calculation.

#### Signature analysis (OncotypeDX and MammaPrint)

The OncotypeDX TCGA cohort included all HR+, HER2-breast cancer patients in the TCGA database who met FDA guideline criteria for the signature, specifically those with 1 to 3 positive axillary lymph nodes or T1b/c-2 pN0 tumors. This resulted in a total of 295 patients with both slide images and survival data available. Similarly, the MammaPrint TCGA cohort included all breast cancer patients in the database who met FDA guideline criteria for the signature, including those with stage I-II disease, tumor sizes ≤ 5.0 cm, and lymph node-negative status. This selection resulted in 367 patients with both slide images and survival data available.

To apply the original OncotypeDX and MammaPrint signatures to TCGA RNA-seq data (**Supplementary Fig. 6a**), scores were calculated using the genefu^67^ R package (version 2.37.0). OncotypeDX recurrence risk groups were classified as high-risk (recurrence score > 30), intermediate-risk (18–30), and low-risk (< 18).

For TCGA model development using RNA-seq measurements and SPAND (**Fig. 5c, Supplementary Fig. 6b,c**), elastic-net regularized Cox proportional hazards survival models were constructed with the lifelines^68^ package (version 0.28.0). Robust performance was ensured through Kaplan-Meier survival analysis and concordance index (C-index) calculations, using five-fold cross-validation repeated 1,000 times. Patient risk scores were obtained by averaging predictions from the validation folds across all iterations.

#### Bivariate SPAND analysis

To investigate the role of spatial co-expression between ER/HER2 and their protein-protein interaction (PPI) partners in patient survival, we calculated the bivariate Moran’s I index between the imputed expression of *ESR1* and its key interacting proteins, identified using STRING^46^. The bivariate Moran’s I was computed using the global_moran_bv function from the sfdep^69^ (0.2.5) package in R, with spatial weights defined as the inverse of the distance, normalized by tile size. To adjust for gene expression magnitude, the bivariate Moran’s I was scaled by multiplying it with the mean imputed bulk expression of the two interacting genes.

#### HER2 SPAND

To estimate HER2 expression in cancer cells, we applied *Path2Space*-inferred spatial gene expression to compute the normalized enrichment score (NES) for the ‘ErbB Signaling Pathway (WP673)’ from the WikiPathways library^48^. NES values were calculated per pseudospot using ssGSEA implemented in GSEAPY. To isolate HER2 activity specifically in cancer cells, NES values were adjusted by dividing by the cancer cell fraction, estimated by the cell type abundance model at each pseudospot. This yielded HER2 expression per cancer cell. SPAND analysis was performed on these normalized values.

### Chemotherapy and trastuzumab response prediction models

#### *Path2Space* cluster model

Cluster proportions were used as input features to predict response. A pipeline was trained consisting of feature standardization using Scikit-learn’s StandardScaler, followed by L1-regularized logistic regression. During cross-validation, StandardScaler was fitted to each training fold and applied to both the training and test folds to ensure consistent scaling. Model scores and labels were aggregated across all test folds to evaluate performance.

For external validation, predictions from the five cross-validation models were averaged to obtain the final prediction score. In cases where clusters from the TransNEO development cohort were absent in the IMPRESS cohort, their values were set to zero to maintain input consistency.

#### *Path2Space* Combined Model for Trastuzumab

The combined model predicted trastuzumab response by averaging the prediction scores from the Cluster Model and HER2 SPAND. SPAND scores were standardized using StandardScaler in the same cross-validation pipeline to ensure consistency. For the IMPRESS cohort, scaling factors (mean and standard deviation) were averaged across the five cross-validation folds. The final prediction score was the average of the cluster model and scaled SPAND score.

#### Bulk Model

To evaluate the predictive power of a model based on inferred bulk gene expression, we developed a control model with a similar architecture to *Path2Space*. This bulk expression prediction model was trained on the BRCA-TCGA cohort (n = 869 slides with matched H&E images and bulk gene expression data) to predict bulk gene expression values from H&E slides.

The model’s architecture mirrors that of *Path2Space*, utilizing the same image preprocessing and feature extraction pipeline. An MLP regressor was trained to predict bulk gene expression for each tile. Predictions were then averaged across all tiles from the same slide to generate inferred bulk expression values for each sample.

Training was conducted using a 5×5 nested cross-validation strategy. The resulting model was applied to the TransNEO and IMPRESS cohorts to infer bulk expression values. For the TransNEO cohort, a logistic regression model was trained to predict response based on the inferred bulk values. The data were first scaled using Scikit-learn’s StandardScaler and the top 11 features were selected via SelectKBest. An L1 penalty was applied to the logistic regression model to enhance feature sparsity. This model was trained in five-fold cross-validation on both trastuzumab-treated and chemotherapy datasets and validated on the IMPRESS cohort.

#### Direct Model

To assess the feasibility of predicting treatment response directly from slide images, we developed a second control model using the same image preprocessing and feature extraction pipeline as *Path2Space*. This direct prediction model bypasses gene expression inference by training an MLP classifier to predict response directly from extracted image features.

For model training, each slide was partitioned into tiles of 500×500 pixels at 20x magnification, with each tile assigned the label of the entire slide. The model was trained using a 5×5 nested cross-validation strategy on the TransNEO cohort and validated on the IMPRESS cohort to evaluate its generalizability to independent datasets.

#### Other methods

We compared our models to previously published models which also predicted trastuzumab and chemotherapy response in TransNEO the IMPRESS cohorts. We obtained the scores for the Sammut-ML model^31^ from transneo-diagnosis-MLscores.tsv downloaded from https://github.com/cclab-brca/neoadjuvant-therapy-response-predictor. We used the scores for the Clinical+DNA+RNA+DigPath model. We collected the scores for the ENLIGHT model^56^ from https://ems.pangeabiomed.com/. For the Huang et al. model^32^, we extracted the AUCs directly from Table 2 in the paper, as the scores were not provided. We did not report the 95% confidence intervals as these were also not available. We used the AUCs for the model based on the H&E only to ensure a fair comparison with our model. Finally, for the models from Aswolinskiy et al.^55^, we obtained the AUCs and 95% confidence intervals directly from Table 4 in the paper. We compared our method to the models based on the ratio of lymphocytes to tumor in the slide (LTR), the ratio of (inflamed) tumor close to lymphocytes to the overall tumor amount (ITR), the computational tumor infiltrating lymphocytes score (cTILs) and the mitotic rate (MTR) as the number of detected mitoses within tumor regions divided by the tumor area.

#### Response model evaluation

For model performance evaluation, we used AUC, AUPRC, as implemented in Scikit-learn, and odds ratio metrics.

To calculate the odds ratio for the *Path2Space*, bulk and direct model, we first use the Youden’s J^57^ statistic to learn optimal thresholds for each of the methods on the training data (TransNEO). For each method, we identified the threshold that maximized J, defined as:

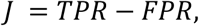

where TPR is the true positive rate and FPR is the false positive rate.

For the test cohorts (IMPRESS), we applied the Haldane-correction^70,71^ to the contingency table to address potentially undefined odds ratio values. This correction involved adding 0.5 to each cell of the contingency table. The corrected odds ratio (OR) was then calculated as:

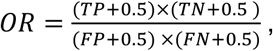

where TP, TN, FP, and FN represent true positives, true negatives, false positives, and false negatives, respectively.

We estimated p-values for each odds ratio using the following steps:

1. Calculate standard error (SE):

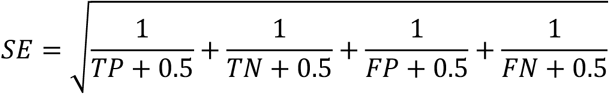
2. Compute test statistic Z: 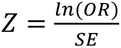
3. Determine p-value: *p* = 2 ⋅ (1 − *Φ*(∣ *Z* ∣))

Where *Φ*(∣ *Z* ∣) is the cumulative distribution function (CDF) of the standard normal distribution.

#### Implementation details

Analyses were performed using Python 3.10.8 and R 4.3.2. Data processing utilized NumPy (1.24.4), Pandas (2.2.2), and scikit-learn (1.5.1). Visualizations were generated with Matplotlib (3.7.2) and Seaborn (0.13.2). Image processing, including tile partitioning and color normalization, was conducted using OpenSlide (1.3.1; for TGGA slides), OpenCV (4.6.0), and Pillow (9.2.0f, or ST slides). Deep learning models were implemented in PyTorch (2.4.0+cu121). Survival analyses were performed with Lifelines (0.28.0) in R. Additional statistical analyses used scikit-learn, and R visualizations were created with ggplot2 (3.5.1).

## Data availability

Spatial transcriptomics gene expression and images from the Bassiouni et al. dataset are available at https://www.ncbi.nlm.nih.gov/geo/query/acc.cgi?acc=GSE210616. The HER2+ dataset can be accessed at https://doi.org/10.5281/zenodo.4751624, and the 10x dataset is available at https://www.10xgenomics.com/datasets. The Janesick et al. dataset is available at https://www.ncbi.nlm.nih.gov/geo/query/acc.cgi?acc=GSM7782699. PanopTIL annotations and images are available at https://sites.google.com/view/panoptils/. H&E-stained whole-slide images and patient response data from the TransNEO-breast dataset are available at https://ega-archive.org/studies/EGAS00001004582, and data from the IMPRESS cohort can be accessed at https://tinyurl.com/IMPRESS-DATA.

## Code availability

*Path2Space* code will be made available for academic research purposes after publication.

