## Supplementary Information for "AI-Driven Spatial Transcriptomics Unlocks Large-Scale Breast Cancer Biomarker Discovery from Histopathology"

### Supplementary Figures

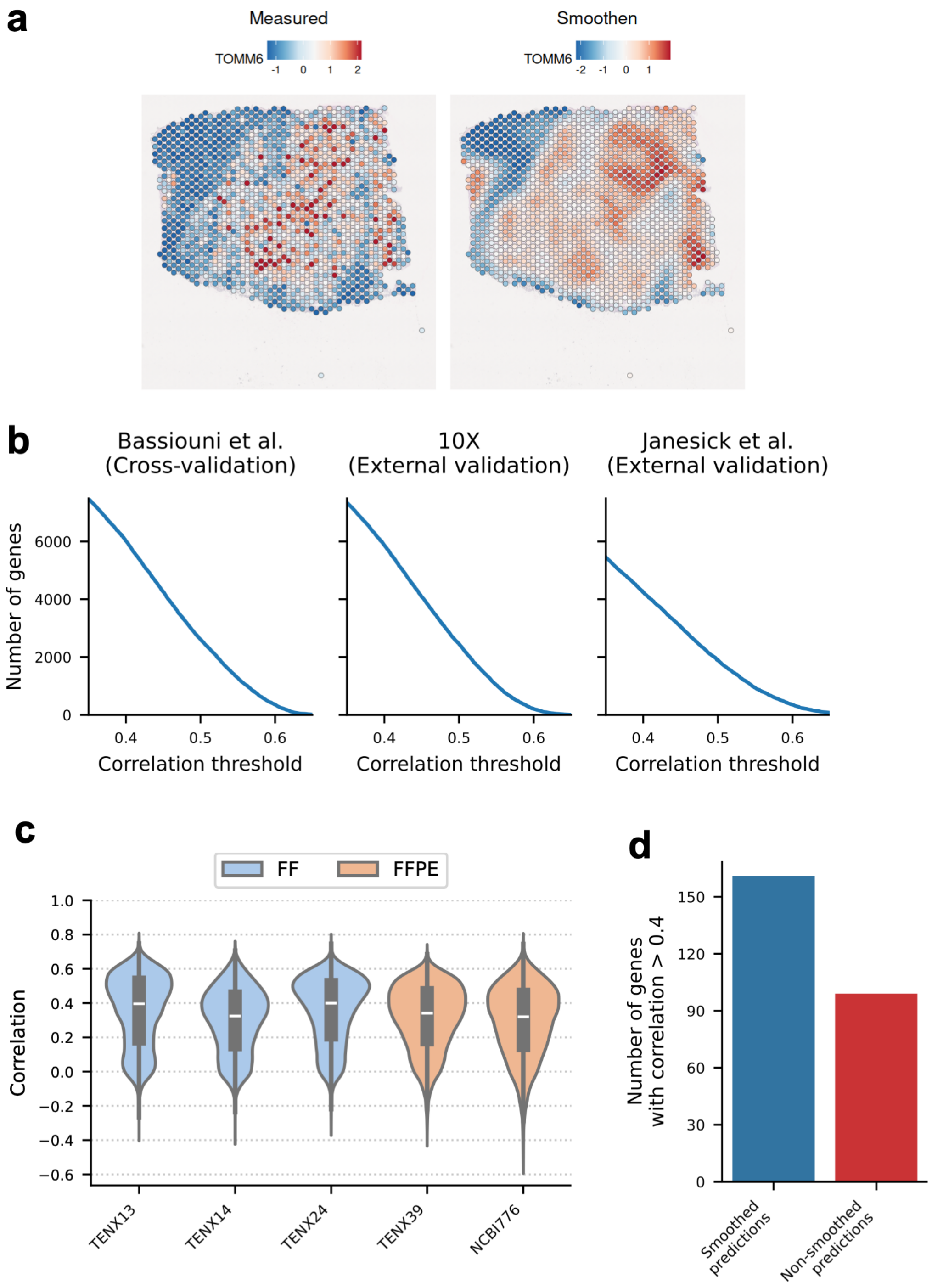

**Supplementary Fig. 1: a,** Visualization of spatial smoothing of gene expression**.** An example slide from the Bassiouni et al. cohort displaying the raw expression of TOMM6 (left) compared to the smoothed expression (right). Colors represent scaled expression levels, with blue indicating low expression and red indicating high expression. **b,** Curves depicting the number of genes surpassing specific correlation thresholds, shown for the Bassiouni et al. cohort (left), 10x cohort (middle), and Janesick et al. (right). **c**, Violin plots of PCC coefficients comparing predicted and actual gene expression (y-axis) across slides (x-axis) for 14,068 genes from the 10X and Janesick et al. cohorts. Blue violins represent fresh frozen (FF) slides, while orange violins represent FFPE slides. The slide NCBI776 is from Janesick et al.; the others are from the 10X cohort. **d,** Boxplot comparing the number of well predicted genes (genes with correlation>0.4) for the smoothed predictions versus the non-smoothed predictions in the legacy spatial transcriptomics cohort used to benchmark against the Hist2ST and HisToGene methods.

1

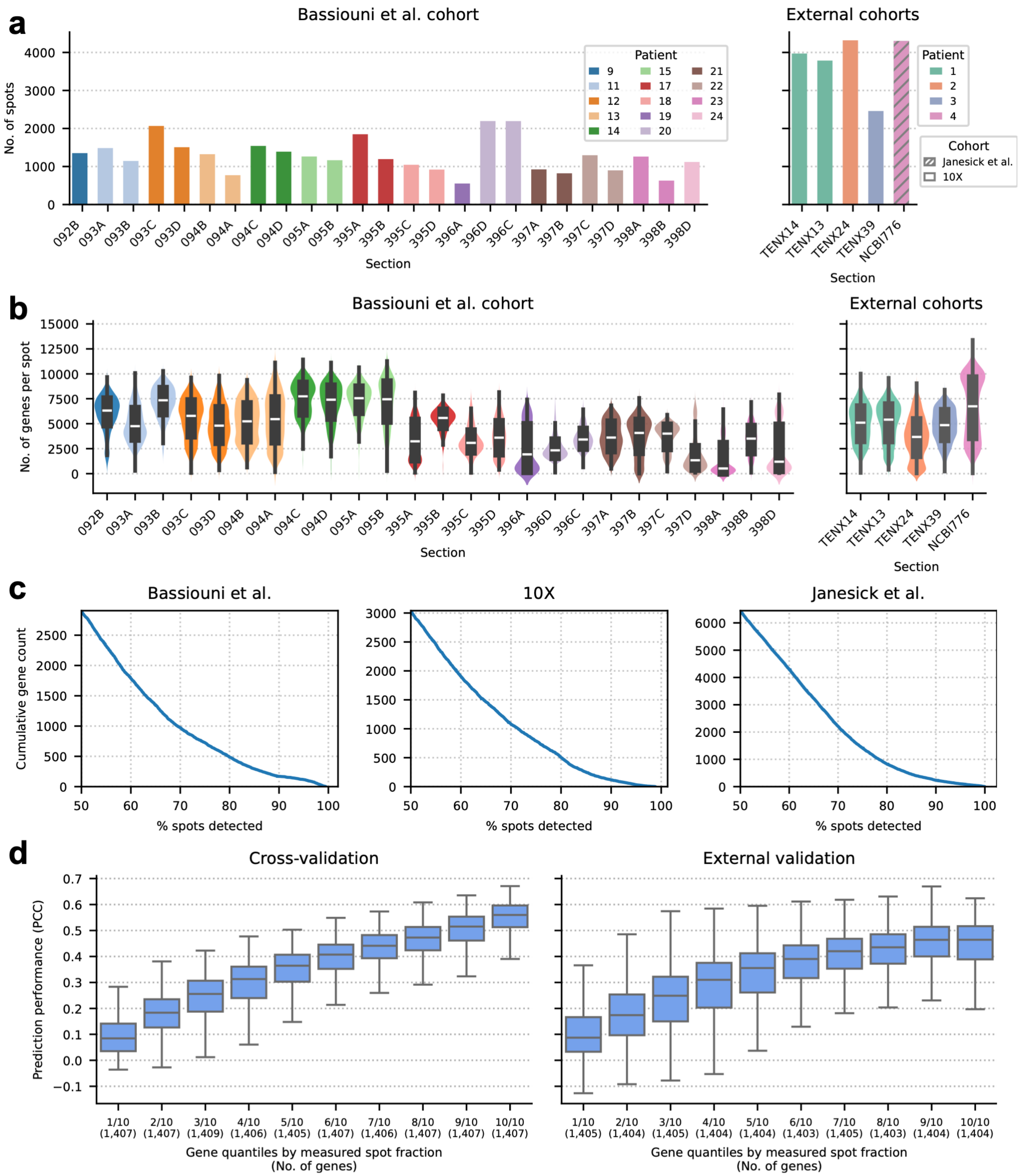


e

| Cohort | Patient | Section | ER | PR | HER2 |
|---|---|---|---|---|---|
| 10X | 1 | TENX13, TENX14 | + | - | + |
| 10X | 2 | TENX24 | + | + | - |
| 10X | 3 | TENX39 | Unknown | Unknown | Unknown |
| Janesick et al. | 4 | NCBI776 | + | - | + |

**Supplementary Fig. 2: Overview of Bassiouni et al. and external validation cohorts with *Path2Space* performance by spot expression. a,** Bar plots showing the number of spots per slide in the Bassiouni et al. dataset, colored by patient, across the Bassiouni et al. (left), 10X, and Janesick et al. (right) cohorts. **b,** Violin plots depicting the distribution of expressed genes per spot for each slide, colored by patient as in panel (**a**). The Bassiouni et al. cohort is on the left, with the 10X and Janesick et al. cohorts on the right. **c,** Cumulative distribution of detected genes across spatial transcriptomic spots. The y-axis represents cumulative gene counts detected in more than a given percentage of spots (x-axis) for each cohort. **d,** Box plots showing the correlation between inferred and actual gene expression across spots. Spots are binned by quantiles based on the fraction of spots with expressed genes. Higher quantiles indicate more expressed (non-zero) genes per spot. Results are shown for cross-validation (Bassiouni et al.) and external validation (mean of 10X and Janesick et al. cohorts). **e,** ER, PR, and HER2 status for patients in the external validation cohorts (10X and Janesick et al.).

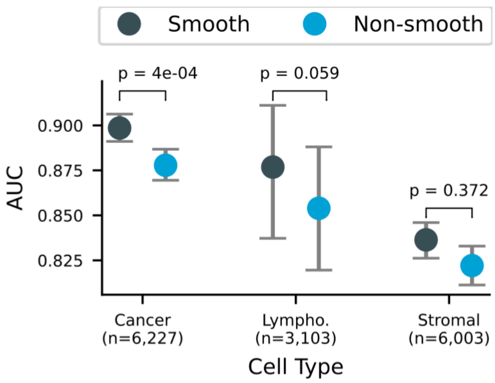


**Supplementary Fig. 3: *Path2Space* deconvolution accuracy comparison using smooth versus non-smooth spatial gene expression.** Area under the receiver operating characteristic curve (AUC) for cell type deconvolution using *Path2Space* with smoothed (black) and non-smoothed (blue) H&E-derived spatial gene expression. Points indicate mean AUC values; error bars show 95% confidence intervals from 1,000 bootstrap iterations. *P* values from the two-sided DeLong's test. Sample sizes (n) are shown for each cell type.

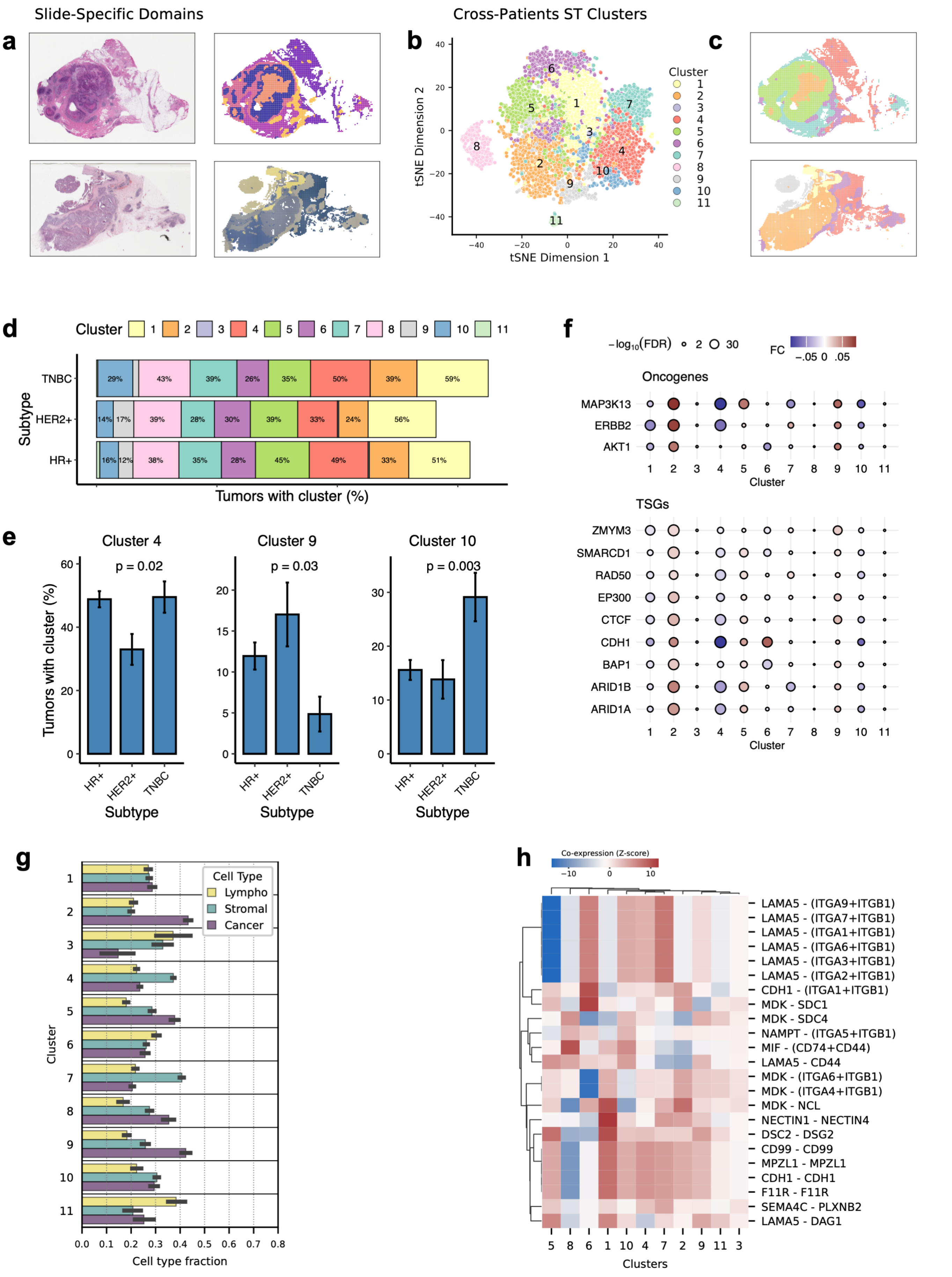

Slide-Specific Domains
Cross-Patients ST Clusters
a
b
c
d
e
f
g
h
tSNE Dimension 1
tSNE Dimension 2
Cluster
Subtype
Tumors with cluster (%)
Cluster 4
Cluster 9
Cluster 10
p = 0.02
p = 0.03
p = 0.003
Oncogenes
TSGs
MAP3K13
ERBB2
AKT1
ZMYM3
SMARCD1
RAD50
EP300
CTCF
CDH1
BAP1
ARID1B
ARID1A
Cell Type
Lympho
Stromal
Cancer
Cell type fraction
Co-expression (Z-score)
Clusters

**Supplementary Fig. 4: Characterization of breast cancer spatial transcriptomic clusters. a**, Examples of slide-specific spatial domains derived using SpaGCN from *Path2Space*-inferred gene expression and spatial organization for two TCGA slides. **b**, t-SNE of domain-averaged pseudospot expression profiles across TCGA slides, identifying 11 distinct cross-patient spatial transcriptomics (ST) clusters. Each point represents a single spatial domain. **c**, TCGA slides colored by their ST cluster assignments from (**b**)**. d,** Percentage of tumors containing each ST cluster across clinical subtypes. **e,** ST clusters showing significant differential representation across clinical subtypes. The y-axis shows the percentage of tumors containing each cluster; error bars represent standard errors. *P* values were calculated from the two-sided chi-squared test. **f**, Dot plot showing the cluster-specific differential expression of breast cancer oncogenes and tumor suppressors across ST clusters. Color indicates average fold change between cluster and other clusters within slides; dot size represents significance (-log FDR) from two-sided linear regression. **g,** Mean fractions of cancer, lymphocyte, and stromal cells across ST clusters. **h,** Heatmap of spatially-resolved ligand-receptor interactions inferred by SpatialDM (likelihood ratio test,FDR<0.05, two-sided). Rows show ligand-receptor pairs, columns represent clusters, and colors indicate global z-scores of Bivariate Moran's I statistics.

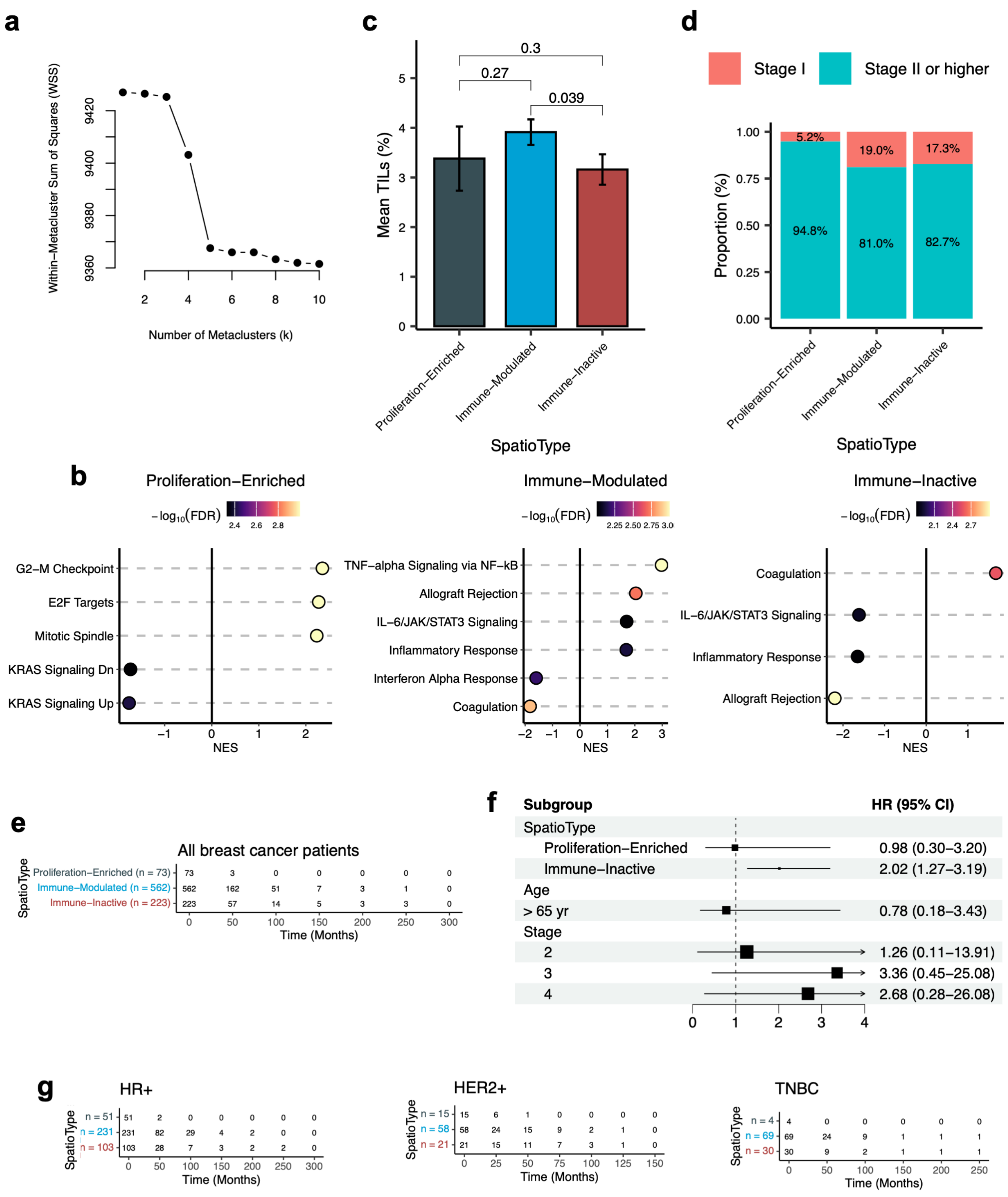


**Supplementary Fig. 5: *SpatioTypes* and their prognostic value. a,** Elbow plot showing the within-meta-cluster sum of squares (WSS) for different numbers of subgroups (*SpatioTypes*) (k).

The plot indicates that three subgroups provide the optimal balance between complexity and explained variance in the TCGA BRCA cohort. **b**, GSEA comparing each *SpatioType* against all others. Rows show cancer hallmark gene sets, with normalized enrichment scores (NES) on the horizontal axis. Color intensity represents -log10 adjusted P values. Shown are significant results (FDR < 0.05) for proliferative hallmarks in the Proliferation-Enriched *SpatioType* and immune hallmarks in the Immune-Inactive and Immune-Modulated *SpatioTypes*. **c**, Mean percentage of TILs (Tumor Infiltrating Lymphocytes) across *SpatioTypes*. *P* values from the single-sided Mann-Whitney U-test. **d,** Percentage of tumors in pathologic Stage I or higher across *SpatioTypes***. h,** Forest plot of multivariate survival analysis adjusting for age and tumor stage, confirming that the Immune-Inactive *SpatioType* is significantly associated with poor prognosis. **i**, Number of patients at risk over time for each *SpatioType* in the Kaplan-Meier analysis shown in **Fig. 3g**. **j**, Number of patients at risk over time for each *SpatioType* within clinical subtypes (HR+ [ER+ or PR+], HER2+, and TNBC) corresponding to **Fig. 3h**. The *SpatioTypes* are color coded as in **Fig. 3g,h** and panel (**i**).

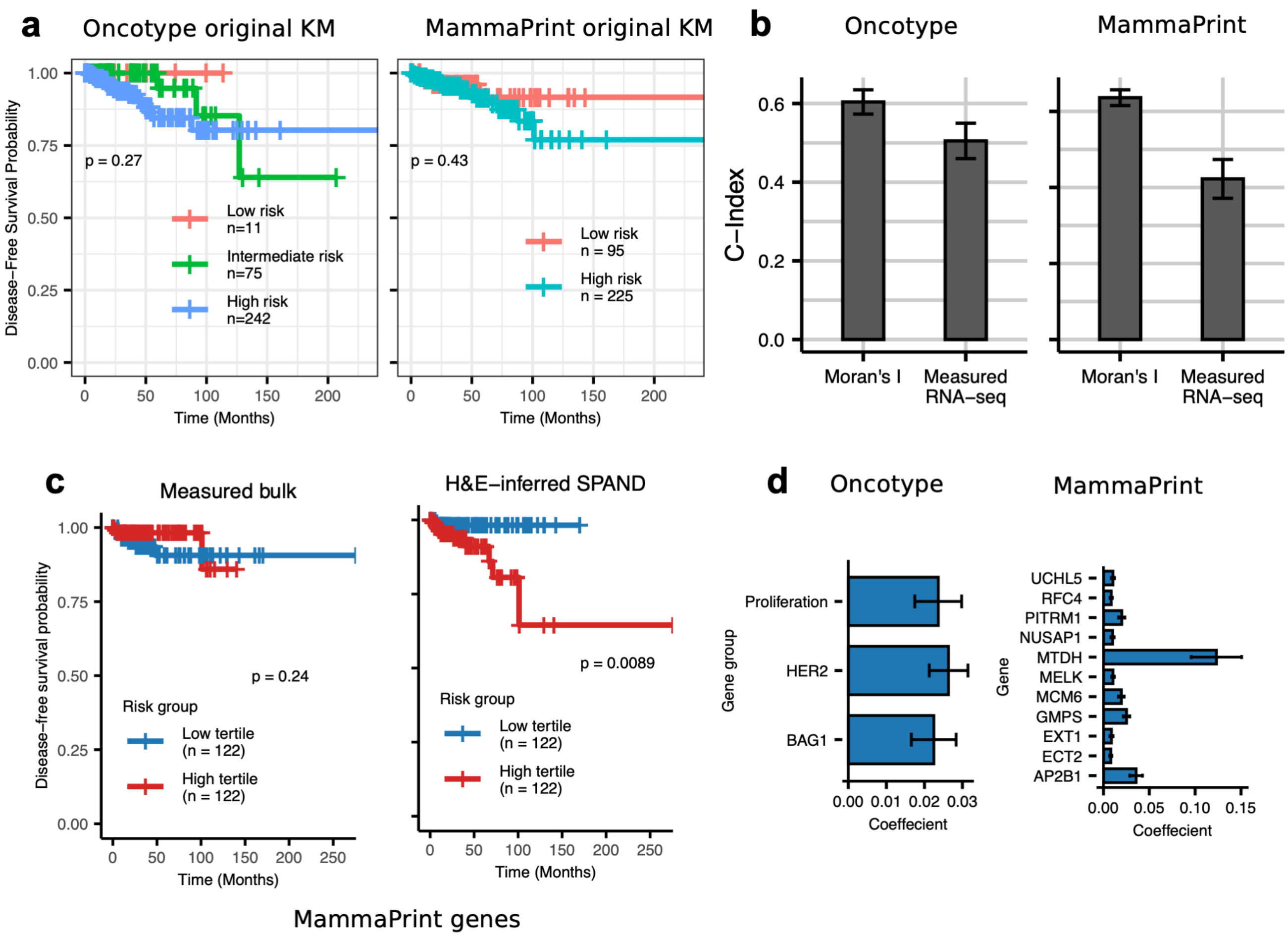

**Supplementary Fig. 6: Validation of SPAND-based breast cancer risk score signatures. a,** Kaplan-Meier curves of original OncotypeDX (left) and MammaPrint (right) signatures applied to TCGA RNA-seq data using the genefu R package. Risk groups assigned by signature criteria; *P* values from two-sided log-rank test. **b,** Cross-validated risk model performance (c-index) comparing bulk RNA-seq and SPAND for OncotypeDX (left) and MammaPrint (right) genes in TCGA. Error bars from 1,000 five-fold cross-validations. **c,** Kaplan-Meier survival curves comparing the predictive power of the *measured* bulk RNA-seq values (left) and the inferred SPAND scores (right) for the MammaPrint (n = 367) signatures in the TCGA. Risk groups show top and bottom tertiles; *P* values from log-rank test. **d,** SPAND model coefficients for OncotypeDX gene groups (left) and MammaPrint genes (right).

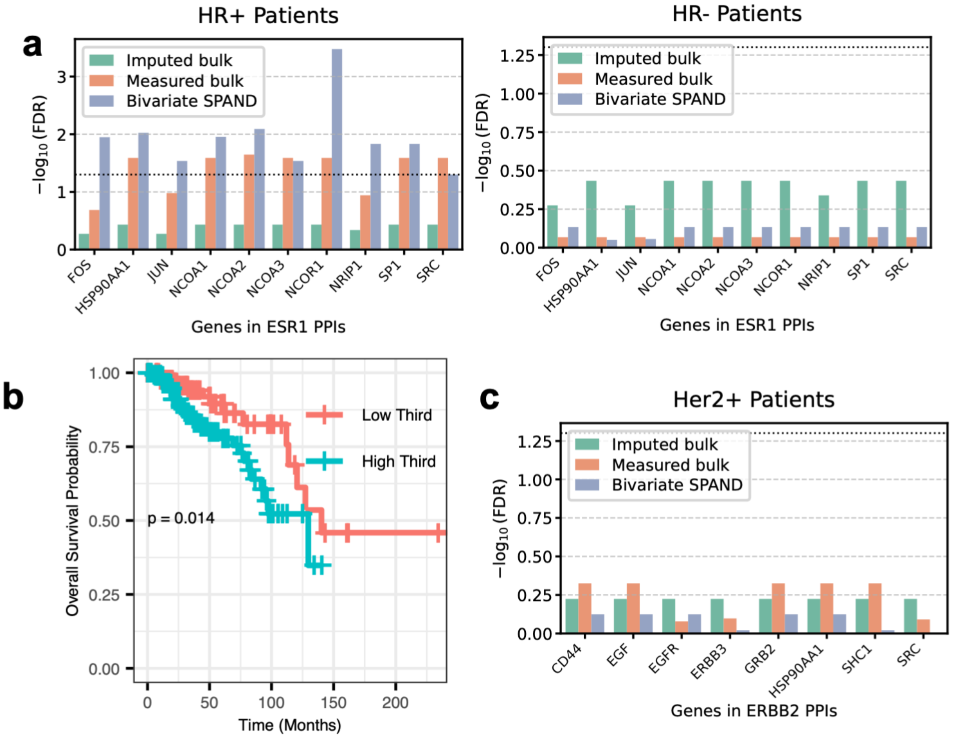


**Supplementary Fig. 7: Association of bivariate SPAND-based scores with overall survival a,** Association with overall survival of *ESR1*-PPI partner pairs in TCGA HR+ (n=385, left) and HR- (n=197, right) breast cancer patients, comparing bivariate SPAND, *Path2Space*-inferred mean expression, and bulk RNA-seq mean expression. Y-axis: -log(FDR) from univariate Cox regression. Related to **Fig. 5d** (HR+ results). **b,** Kaplan-Meier curves for ESR1-NCOR1 PPI pair bivariate SPAND. Risk groups show top and bottom tertiles; *P* values from two-sided log-rank test. **c,** Association with overall survival of *ERBB2*-PPI partner pairs in TCGA HER2-positive breast cancer patients (n=94), comparing bivariate SPAND, *Path2Space*-inferred mean expression, and bulk RNA-seq mean expression. Y-axis: -log(FDR) from univariate Cox regression.

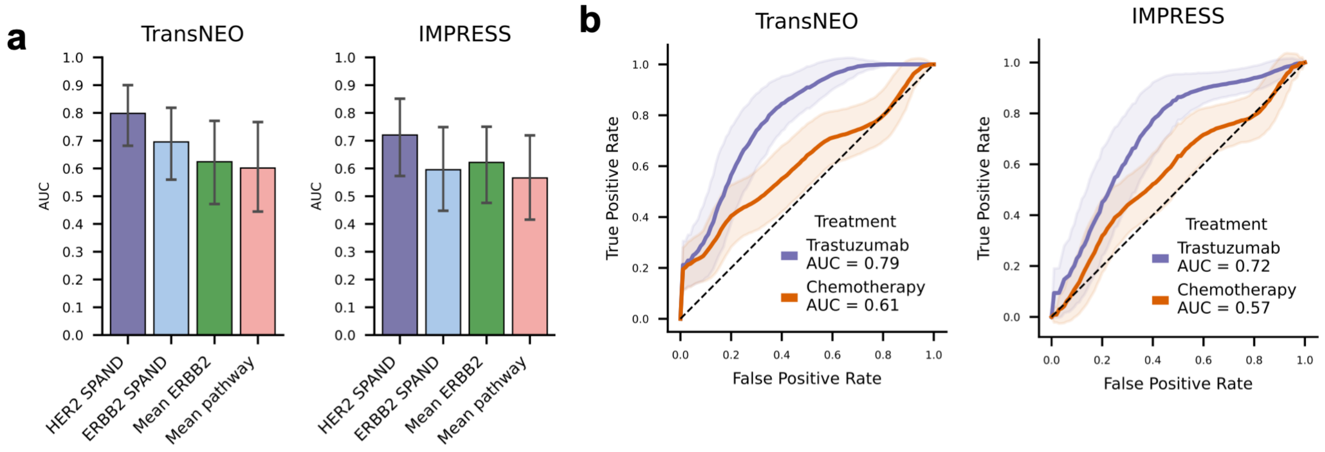


**Supplementary Fig. 8: Validation of HER2 SPAND scores for trastuzumab response prediction. a,** Prediction of Trastuzumab response using H&E-inferred gene expression. Comparison of predictive performance AUCs: HER2 SPAND, *ERBB2* SPAND, mean predicted ERBB2 expression, and mean *ERBB2* pathway activity, all normalized by cancer cell fraction in the TransNEO cohort and IMPRESS cohort. Error bars represent the 95% confidence intervals computed over 1,000 bootstrap iterations. **b,** ROC curves for HER2 SPAND within cancer cells in predicting therapy response. Left: trastuzumab-treated versus chemotherapy-treated patients in TransNEO and IMPRESS cohorts. Right: comparison of H&E-inferred SPAND, mean expression, and Moran's I in trastuzumab-treated patients. Shaded areas: bootstrap standard errors (n=1,000); AUC values in legend.

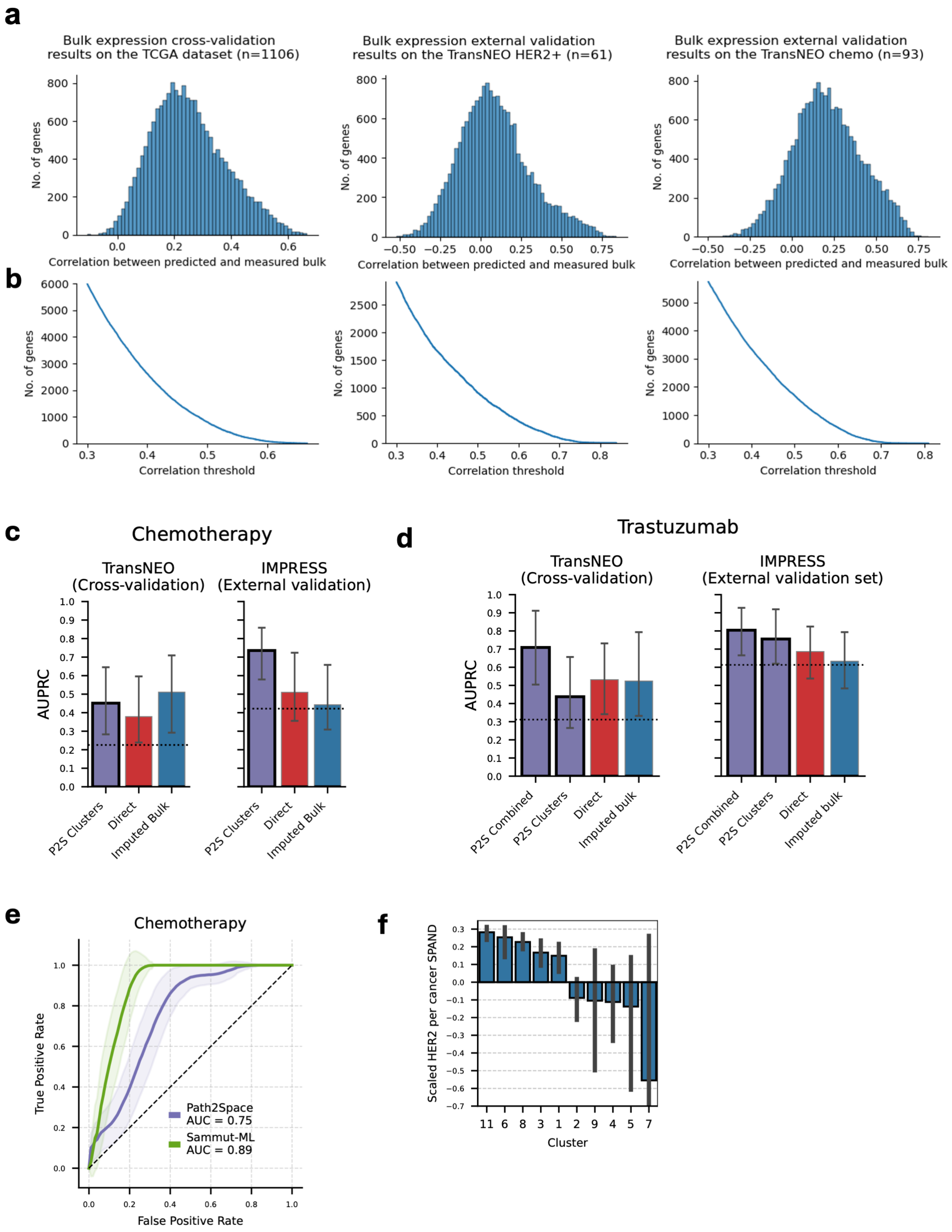
a
Bulk expression cross-validation results on the TCGA dataset (n=1106)
Bulk expression external validation results on the TransNEO HER2+ (n=61)
Bulk expression external validation results on the TransNEO chemo (n=93)
No. of genes
Correlation between predicted and measured bulk
b
No. of genes
Correlation threshold
c
Chemotherapy
TransNEO (Cross-validation)
IMPRESS (External validation)
AUPRC
P2S Clusters
Direct
Imputed Bulk
d
Trastuzumab
TransNEO (Cross-validation)
IMPRESS (External validation set)
AUPRC
P2S Combined
P2S Clusters
Direct
Imputed bulk
e
Chemotherapy
True Positive Rate
False Positive Rate
Path2Space AUC = 0.75
Sammut-ML AUC = 0.89
f
Scaled HER2 per cancer SPAND
Cluster
11 6 8 3 1 2 9 4 5 7

**Supplementary Fig. 9: Evaluation of *Path2Space* predictions and comparison with alternative models. a,b,** Assessment of H&E-based bulk gene expression predictions used in treatment response models compared to *Path2Space*. **a,** Distribution of gene expression correlations in 5-fold nested cross-validation on TCGA (median correlation=0.25, left), and external validation on TransNEO HER2+ (median correlation=0.069, middle) and HER2- (median correlation=0.199, right) cohorts. **b,** Number of genes exceeding correlation thresholds in TCGA (2,615 genes > 0.4, left), TransNEO HER2+ (1,662 genes > 0.4, middle) and HER2- (3,348 genes > 0.4, right) cohorts. **c,d,** Area under the precision-recall curve (AUPRC) for response prediction in TransNEO (cross-validation, left) and IMPRESS (external validation, right) cohorts for chemotherapy (**c**) and trastuzumab (**d**), comparing *Path2Space* with other H&E-based models. *Path2Space* models (thick bar edge): spatial cluster model for both treatments, and combined model (integrating spatial clusters with HER2 per cancer SPAND) for trastuzumab. For IMPRESS, showing Huang et al.'s cross-validation performance and Aswolinskiy et al.'s H&E-based models: lymphocyte-tumor ratio (LTR), computational tumor infiltrating lymphocytes score (cTILs), inflamed tumor ratio (ITR) (orange), and mitoses-tumor ratio (MTR). Error bars: 95% confidence intervals from 1,000 bootstrap iterations. Bar height shows observed AUPRC values. Dashed black horizontal lines indicate overall response rate. **e,** ROC curves comparing *Path2Space* to Sammut-ML (incorporating RNA-seq, genomics, and clinical data) in the TransNEO chemotherapy cohort. Shaded areas: bootstrap standard errors (n=1,000); AUC values in legend. **f,** Mean HER2 SPAND scores for each of the clusters in trastuzumab-treated patients from both cohorts. Error bars denote the standard error.
